# Regulatory Transposable Elements in the Encyclopedia of DNA Elements

**DOI:** 10.1101/2023.09.05.556380

**Authors:** Alan Y. Du, Jason D. Chobirko, Xiaoyu Zhuo, Cédric Feschotte, Ting Wang

## Abstract

Transposable elements (TEs) make up about half of the human genome and many have the biochemical hallmarks of tissue- or cell type-specific *cis*-regulatory elements. While some TEs have been rigorously documented to contribute directly to host gene regulation, we still have a very partial view of their regulatory landscape. Leveraging Phase 4 ENCODE data, we carried out the most comprehensive study to date of TE contributions to the regulatory genome. Here we investigated the sequence origins of candidate *cis*-regulatory elements (cCREs), showing that ∼25% of human cCREs comprising 236,181 elements are derived from TEs. Human-mouse comparisons indicate that over 90% of TE-derived cCREs are lineage-specific, accounting for 8-36% of lineage-specific cCREs across cCRE types. Next, we found that cCRE-associated transcription factor (TF) binding motifs in TEs originated from TE ancestral sequences significantly more than expected in all TE classes except for SINEs. Using both cCRE and TF binding data, we discovered that TEs providing cCREs and TF binding sites are closer in genomic distance to non-TE sites compared to other TEs, suggesting that TE integration site influences their later co-option as regulatory elements. We show that TEs have promoted TF binding site turnover events since human-mouse divergence, accounting for 3-56% of turnover events across 30 TFs examined. Finally, we demonstrate that TE-derived cCREs share similar features with non-TE cCREs, including massively parallel reporter assay activity and GWAS variant enrichment. Overall, our results substantiate the notion that TEs have played an important role in shaping the human regulatory genome.

---

Barbara McClintock, who discovered TEs in maize^1^, was the first to recognize their ability to act as *cis*-regulatory elements (CREs) controlling the expression of nearby genes. In the ensuing decades, it became clear that a large fraction of the genome of multicellular organisms consists of interspersed repeats primarily derived from TEs. In mammals, TEs account for 28-75% of the genome sequence^2^. In humans, at least 46% of the ∼3.1GB haploid genome is derived from TEs^3^. Most TEs in the human genome can be classified into LINE, SINE, LTR and DNA transposon classes. LINEs are autonomous retrotransposons that use target primed reverse transcription to insert into the genome. SINEs are short, non-autonomous elements that rely on the LINE machinery to mobilize. LTR elements in the human genome are mostly derived from endogenous retroviruses (ERVs) which expand using the retroviral replication mechanism. Unlike the other three classes which use RNA intermediates to transpose, DNA transposons mobilize directly via a “cut-and-paste” DNA mechanism^4^. The vast majority of human TEs have long ceased transposition activity, and only a small subset of LINEs and SINEs are known to be currently capable of mobilization in modern humans.

Although McClintock viewed TEs as essential “controlling elements”, it is clear that the vast majority of TE sequences in the human genome have not evolved under functional constraint and therefore do not appear to contribute significantly to organismal fitness^5^. Still, about 11% of evolutionarily constrained bases in human fall under TEs^5^, and there have been many reports showing that some TEs function as CREs, including promoters and enhancers, regulating important biological processes (reviewed in ^6–11^). However, we still lack a global picture of how many CREs are derived from TEs and how many are truly functional.

Another fundamental question concerns the evolution of TEs from selfish elements to CREs co-opted for gene regulation. One model postulates that TEs ancestrally harbor CREs and transcription factor binding sites (TFBSs) in order to regulate their own genes, which are then occasionally co-opted for regulating host gene expression. Many examples of previously characterized TE-derived CREs are consistent with this ancestral origin model^12–15^. An alternative model is that TEs acquire TFBSs and *cis*-regulatory activity post-insertion through mutation over time. This has been observed for P53, PAX-6, and MYC TFBSs in human Alu SINE elements and circadian clock TFBSs in mouse RSINE1 elements, in which imperfect binding motifs matured into canonical binding motifs over evolutionary time^16–18^.

The ENCODE and Roadmap projects have sought to characterize the landscape of CREs in the human genome, providing invaluable resources for scientists all over the world^19,20^. Data from these projects have facilitated systematic investigation of TE contributions to regulatory functions in the genome^21,22^. In ENCODE phase 3, genome-wide annotations of candidate *cis*- regulatory elements (cCREs) were created in both human and mouse genomes^19^. Based on four epigenomic assays and gene annotation, cCREs were classified into promoter-like sequence (PLS), proximal enhancer-like sequence (pELS), distal enhancer-like sequence (dELS), high-H3K4me3 elements (DNase-H3K4me3), and potential boundary elements (CTCF-only). Regions with enhancer signal were separated into pELS and dELS based on their distance to annotated transcription state sites (TSSs). DNase-H3K4me3 cCREs represent regions with promoter signal without a nearby annotated TSS. CTCF-only cCREs represent regions that could be genome folding anchors or other architectural elements. Altogether, cCREs comprise 7.9% and 3.4% of human and mouse genomes, respectively. In the latest ENCODE phase 4, functional assays such as massively parallel reporter assay (MPRA) have been included to validate regulatory element predictions. Here we leverage these resources to quantify the contribution of TEs to the regulatory genome and derive general principles for how TEs become regulatory elements.

## Landscape of TE-derived cCREs in human

To broadly characterize the contribution of TEs to the human regulatory genome, we intersected TEs with cCREs from the version 2 registry of cCREs^19^. As a conservative estimate, we considered cCREs with at least 50% of their sequences coming from a single annotated TE to be TE-derived. Using this criterion, we found that TEs supply ∼25% (236,181/926,535) of all human cCREs (Fig. 1A). When cCREs are separated by annotation type, TE contribution ranges from 4.6% in PLS to 38.2% in CTCF-only cCREs. Compared to their genomic proportion, TEs are generally underrepresented in all types of cCREs (Extended Data Fig. 1). Notably, TEs are most depleted in PLS, possibly due to a combination of purifying selection against TE insertion in promoters and incomplete annotation of TE promoters. By contrast, DNase-H3K4me3 and CTCF-only cCREs were enriched for the LTR class of elements (log_2_ enrichments of 0.42 and 0.46, respectively). These results suggest that LTR elements have been a prominent source of non-canonical promoters and CTCF binding sites, an observation consistent with previous reports^23–26^.

**Fig. 1:**
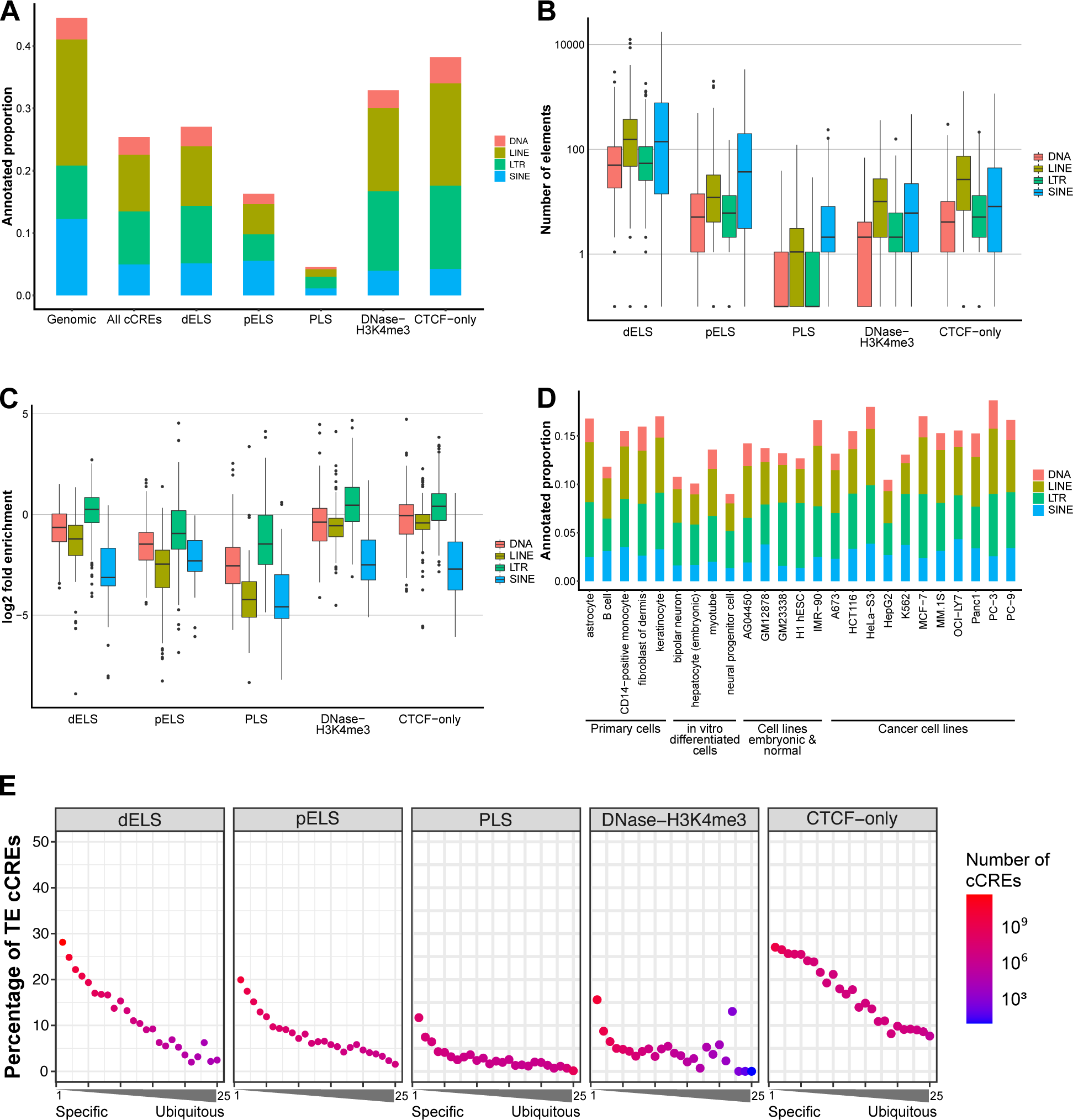
Overlap of TEs with human cCREs. A) Proportion of genome and cCREs that are TE-derived. B) Number of elements per TE subfamily, grouped by TE class, that are associated with a cCRE. C) Enrichment of TE subfamily overlapping with cCREs relative to their abundance in the genome, grouped by TE class. D) Proportion of cCREs that are TE-derived across 25 fully profiled cell/tissue types. E) Percentage of cCREs that are TE-derived for cell/tissue specific cCREs to ubiquitously used cCREs. The x-axis is the number of fully profiled cell/tissue types in which the cCRE is found. For all boxplots in this paper, box, interquartile range (IQR); center, median; whiskers, 1.5 × IQR.

Within each TE class, TEs are further subdivided into many subfamilies of variable age and sequence composition. Thus, we next examined TE contributions to the human regulatory genome at the subfamily level. In terms of absolute numbers of cCRE-associated TEs, LINE and SINE classes contribute the most cCREs per subfamily on average (Fig. 1B). On the other hand, after normalizing to genomic abundance, the LTR class is the most enriched per subfamily on average for cCREs (Fig. 1C). These results confirm that LTR elements are generally more likely to supply cCREs in the human genome, possibly because they contain strong promoter and enhancer sequences^10^. However, numerically the majority of TE-derived cCREs come from SINEs and LINEs due to their sheer abundance in the human genome.

Considering that regulatory elements are often active in a cell-type specific manner, we evaluated the contribution of TEs to each of the 25 ENCODE cell/tissue types with full cCRE profiling. Overall, TEs make up between 9-19% of cCREs across cell/tissue types (Fig. 1D). The proportion of TE classes contributing to cCREs stays relatively stable across cell/tissue types (Extended Data Fig. 1). Next, we examined whether TE-derived cCREs are more or less likely to be cell-type specific compared to non-TE cCREs. We grouped all cCREs by the number of cell types that share them. With more ubiquitously active cCREs across the 25 fully profiled cell/tissue types (i.e. less cell-type specific), the percentage of cCREs that are TE-derived decreases (Fig. 1E), indicating that cCREs contributed by TEs are more likely to be cell-type specific. This observation is consistent with previous reports that find TEs to contribute cell-type specific regulatory elements^21,27,28^.

## Evolution of TE-derived cCREs across human and mouse

Next, we investigated the contribution of TEs in the evolution of cCREs in the human and mouse lineages. Starting from 926,535 human cCREs, we identified syntenic mouse regions using UCSC liftOver^29^, yielding 601,136 syntenic regions corresponding to ∼66% rate of synteny (Fig. 2A, Extended Data Fig. 2). This is significantly higher than the ∼40% rate of synteny based on whole genome comparison (p=1.5×10^−323^, binomial test), which is expected as cCREs should be enriched for functional regulatory elements and therefore more evolutionarily constrained^30^. To identify cCREs derived from TEs orthologous in human and mouse (acquired from their common ancestor), we required that the human cCRE be TE-associated and the corresponding mouse syntenic region contains the same annotated TE (Methods). As expected, orthologous TEs are primarily composed of old TE subfamilies that exist in both human and mouse (Extended Data Fig. 2). This approach identified 18,010 TE-derived human cCREs (1.9% of all human cCREs) with a mouse orthologous sequence. Overall, 97% (228,670/236,181) of human TE-derived cCREs are only found in the human lineage. We performed the reciprocal analysis starting from mouse cCREs and found similar results: 1.7% (5,900/339,815) of mouse cCREs are TE-derived and have human orthology, and 93% (38,815/41,800) of mouse TE-derived cCREs are only found in the mouse lineage. Thus, TE-derived cCREs are overwhelmingly lineage-specific, and ancient TEs are a minor source of cCREs shared between human and mouse.

**Fig. 2:**
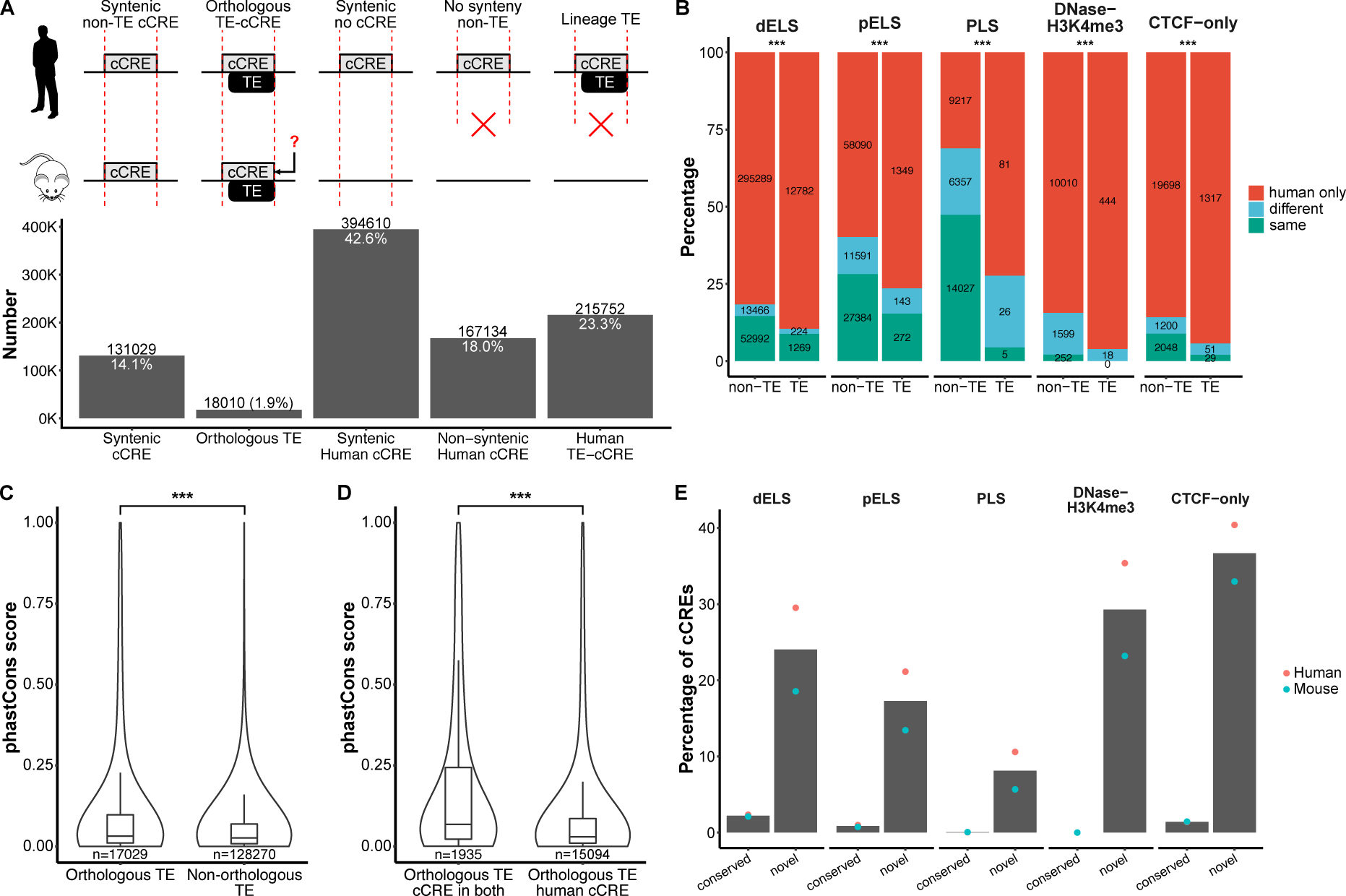
TE-derived conserved and lineage-specific cCREs in human and mouse. A) Classification of shared and lineage-specific cCREs for human to mouse comparison. For orthologous TE-cCREs, syntenic cCRE in mouse is not required but can be present. B) Percentage of cCREs that are shared or lineage-specific for orthologous TE and syntenic non-TE human anchored cCRE regions. Shared cCREs are split into “same” and “different” categories depending on the syntenic human and mouse cCRE types. Grouping by cCRE type is done using the anchored human cCRE. C) 100-way vertebrate phastCons score distributions for orthologous TEs and non-orthologous TEs associated with human cCREs. D) 100-way vertebrate phastCons score distributions for orthologous TEs that have cCRE in both human and mouse vs. human only. E) Percentage of conserved and novel (lineage-specific) cCREs that are TE-derived, split up by cCRE type. Percentages for human and mouse are shown by red and blue dots, respectively. Bars represent the mean percentage between human and mouse. *** p < 0.001

To investigate how often orthologous TEs evolve shared function across lineages, we first searched for orthologous TEs with conserved *cis*-regulatory function in human and mouse. Of 98,278 human cCREs with the same syntenic mouse cCRE, 1,575 (1.6%) are derived from orthologous TEs. This is similar to the percentage of human cCREs that are TE-derived and have a mouse TE ortholog (1.9%), indicating that conserved regulatory elements are not enriched for TEs. We next asked whether orthologous TEs and non-TE syntenic sequences have different annotations between human and mouse. We categorized each human-mouse pair of sequences into same cCRE type (same), shared cCREs but different type (different), or lineage-specific cCREs. Regardless of cCRE type, orthologous TEs contributing cCREs in human display a significantly different proportion of same, different, and lineage-specific cCRE annotations compared to non-TE sequences (Fig. 2B). Contrary to the null expectation where the proportions are the same between TEs and non-TEs, orthologous TEs that contribute cCREs are more lineage-specific than the non-TE syntenic background, ranging from 7.9% difference for dELS to 41.2% difference for PLS in human (Exact multinomial tests, p<0.001). We performed the same analyses starting from mouse cCREs and found similar results, with differences ranging from 8.7% for DNase-H3K4me3 to 36% for PLS (Extended Data Fig. 2, exact multinomial tests, p<0.001). This suggests that among cCREs with a shared sequence origin, TEs are more likely to diverge in *cis*-regulatory activity to provide lineage-specific function compared to non-TE sequences.

Sequence conservation is generally a good indicator for conserved function. Since we can be confident that orthologous TEs in human and mouse descend from a common ancestor, we tested whether sequence conservation as measured by phastCons score is correlated with their shared annotation as cCREs. Considering only TE subfamilies that are ancestral to human and mouse, we confirmed that orthologous TE sequences have higher phastCons scores than non-orthologous TEs (Wilcoxon test, p < 2.2×10^−16^) (Fig. 2C). Furthermore, we found that orthologous TEs with shared cCRE annotation have higher phastCons scores compared to orthologous TEs with lineage-specific cCRE annotation (Wilcoxon test, p < 2.2×10^−16^) (Fig. 2D). This result suggests that TE-derived cCREs shared by both species are under stronger functional constraint than those that are lineage-specific cCREs. Thus, this set of ∼1,500 orthologous TE-derived cCREs are strong candidates for being co-opted for important and conserved cellular function.

Given that most human TE-cCREs are not found in mouse and vice versa, we sought to quantify the contribution of TEs to lineage-specific cCREs relative to non-TE sequences. In human, 85% (788,108/926,535) of cCREs were identified as lineage-specific due to either lack of syntenic sequence in mouse or synteny with no mouse cCRE. Among human lineage cCREs, 29% (228,670/788,108) could be attributed to TEs. In mouse, 61.6% (209,338/339,815) of cCREs were identified as lineage-specific, of which 18.5% (38,815) were TE-associated. We found that TEs have contributed between 6-38% of lineage-specific cCREs depending on cCRE type, with the lowest contribution to PLS and the highest to CTCF binding sites (Fig. 2E). Despite more cCRE data being available for human compared to mouse, we observed a similar trend in human and mouse in which TEs supplied 10-40% of human lineage cCREs and 6-33% of mouse lineage cCREs. Overall, these results support the long-standing hypothesis that TEs have had a substantial impact on *cis*-regulatory innovation during mammalian evolution^31–34^.

## Origin of cCRE-associated transcription factor motifs in TEs

As TFBSs are a major component in driving *cis*-regulatory activity of a sequence, we looked for TF motifs that are associated with cCRE activity in TEs. For each TE subfamily defined by RepeatMasker, we looked for TF motifs that are enriched in cCRE-associated copies of the subfamily relative to non-cCRE copies of the same subfamily (Extended Data Fig. 3A, Methods). By using copies of the same TE subfamily as background sequences in this analysis, we minimize the influence of TF motifs that are merely enriched in the TE subfamily compared to the rest of the genome. In total, we could detect 1183 cCRE-associated TF motifs across 376 TE subfamilies (Extended Data Fig. 3B).

We investigated whether cCRE-enriched TF motifs likely originated from the ancestral TE or arose through mutations after insertion. We first asked what percentage of cCRE-enriched motifs can be identified in the TE’s consensus sequence, which represents their ancestral TE sequence. Of 1183 motifs, 541 (46%) are found in consensus sequences (Extended Data Fig. 3C). Notably, SINEs have the lowest percentage (mean of 12%) among TE classes. To increase resolution and specificity, we extended our analysis to consider motif location for individual TE copies. If a TF motif is truly derived from its ancestral TE insertion, we expect the motif to be in the same relative position as the consensus sequence’s motif. Thus, we inferred the ancestral origin of each TE’s motifs based on the presence or absence of the motif within 10bp of a consensus motif (Fig. 3A). At a mean of 7%, SINEs once again have the lowest percentage of ancestrally derived TF motifs (percent ancestral origin) for cCRE-associated motifs across TE subfamilies (Fig. 3B). We also observed that cCRE-associated TF motifs have significantly higher percent ancestral origin compared to randomly selected motifs for DNA transposons (p = 1.2×10^−8^), LINEs (p = 8.7×10^−8^), LTRs (p < 2.2×10^−16^), and ERV internal regions (ERV-int) (p = 0.0067). This suggests that ancestral TE sequences serve as an important source of TF motifs in cCREs for most but not all TE classes. For SINEs, the percentage of cCRE-associated motifs of ancestral origin was not different from random expectation (p = 0.86), indicating that the ancestral TE sequence does not generally explain the presence of TF motifs enriched within SINE-derived cCREs.

**Fig. 3:**
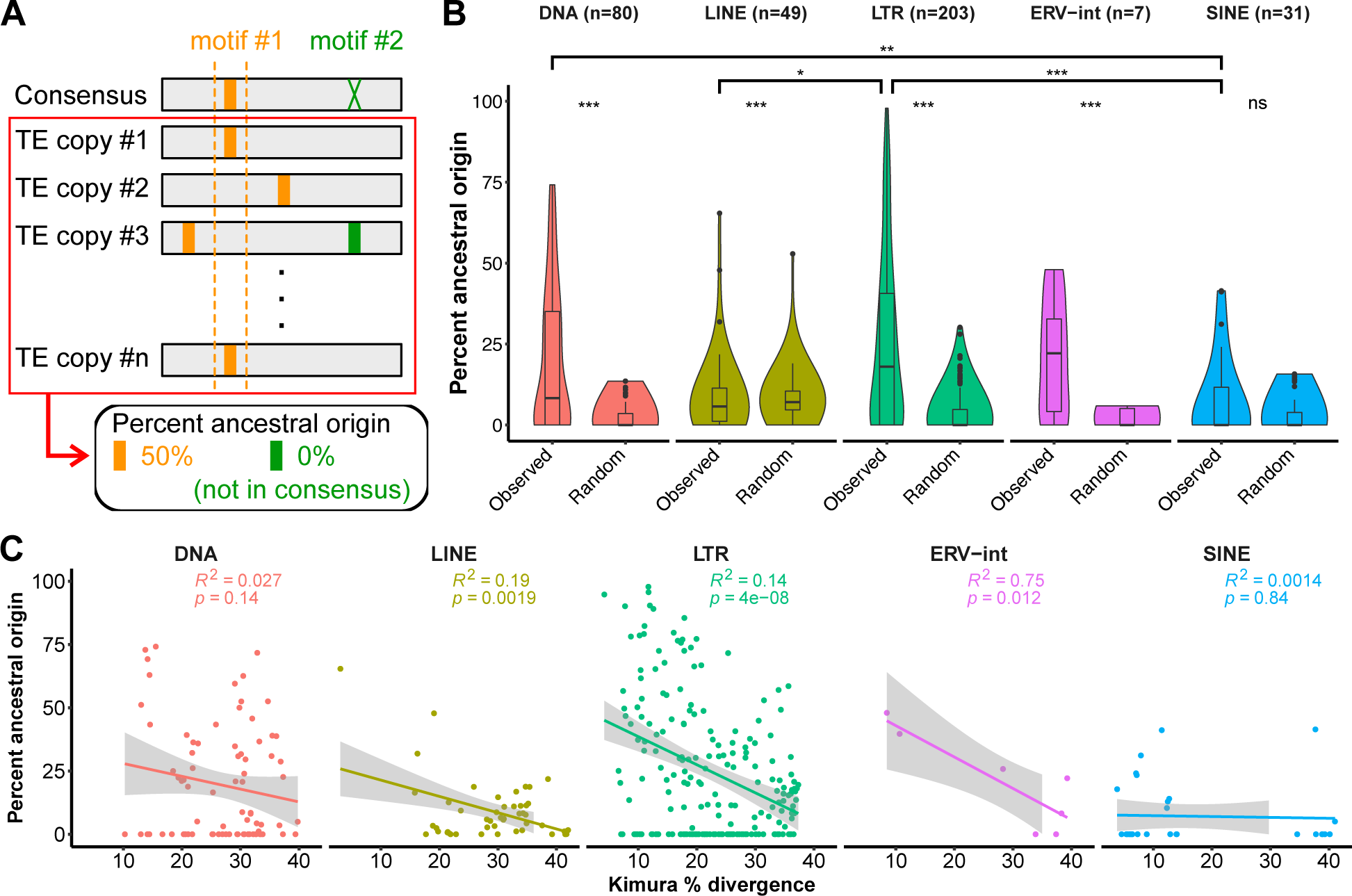
cCRE enriched TF motifs are mostly ancestral except in SINE. A) Analysis workflow for percent ancestral origin calculation of cCRE-associated TF motifs. B) Distribution of mean percent ancestral origin of cCRE-associated TF motif for each TE subfamily, separated by TE class. Two-sided Wilcoxon rank sum test with Benjamini-Hochberg correction was performed to compare percent ancestral origin between observed cCRE associated TF motifs and randomly selected TF motifs, and to compare percent ancestral origin between TE classes. C) Correlation between TE subfamily Kimura divergence and cCRE-associated TF motif percent ancestral origin. R-squared and p-values for each linear regression is shown. * p < 0.05, ** p < 0.01, *** p < 0.001

Next, we considered the evolutionary fate of TF motifs within TE sequences. If TEs contain ancestral TF motifs that are occasionally retained within cCREs, we expect many ancestrally derived TF motifs to gradually decay away through accumulated mutations as most TE copies neutrally evolve. Consistent with this prediction, TE subfamily age, as measured by the mean Kimura divergence of individual copies from the subfamily consensus, is negatively correlated with the percentage of ancestrally derived TF motifs (Fig. 3C). When we break down this analysis per TE class, we found that this correlation held for LINEs, LTRs, and ERV-int, but not for DNA transposons or SINEs. As an orthogonal analysis, we calculated the percentage of TE copies containing the TF motif within each subfamily, based on the hypothesis that ancestrally derived motifs should be found in a higher percentage of copies compared to mutation-derived motifs. TE subfamily age was negatively correlated with the percentage of copies with motif for all TE classes except for SINEs (Extended Data Fig. 3E). Taken together, these findings suggest that most TE subfamilies arrive in the genome already containing *cis*-regulatory sequence features that are retained within cCREs. SINEs tend to adopt a different trajectory whereby TF motifs do not preexist within their ancestral sequence but evolve subsequently via mutations. It is important to note the considerable variation between different TE subfamilies, highlighting that each TE subfamily has its own unique evolutionary path to maintain or acquire *cis*-regulatory activity.

Examining the consensus coverage of cCRE-associated TEs compared to non-cCRE TEs revealed an unexpected enrichment over the 5’ end of LINE1, even after controlling for length (Extended Data Fig. 4 and 5). This indicates that LINE1s that contain the 5’ end disproportionately contribute to cCREs. Our observation is consistent with the prediction that the 5’ region of LINE1 harbors their promoter sequence and contains a wealth of TFBSs^35–38^. These results suggest that the 5’ end of LINEs may be similar to LTRs in providing regulatory sequence.

## Genomic context influences the *cis*-regulatory potential of TEs

As TEs are not evenly distributed throughout the genome, we next sought to explore whether there is any relationship between the genomic loci of TE-derived cCREs and non-TE cCREs. Specifically, we quantified the relative distance from either TEs or cCRE-associated TEs to the nearest non-TE-derived cCREs. If TEs randomly develop into cCREs regardless of their insertion location, we should observe a uniform distribution of cCRE-associated TEs relative to non-TE cCREs. As expected, TE insertions are uniformly distributed in the genome relative to cCREs (red line in Fig. 4A). However, TEs associated with PLS, pELS, and DNase-H3K4me3 are significantly closer to other cCREs of the same category when using a cell type-agnostic approach by Kolmogorov-Smirnov test (KS test) (blue line in Fig. 4A). While not significantly closer when considering cell type-agnostic annotations of dELS, TEs associated with dELS are significantly closer to non-TE dELS sites after separating dELS by cell or tissue type (green line in Fig. 4A). This suggests that, despite being uniformly distributed in the genome in general, TE insertions close to other promoters or enhancers are more likely to be promoters or enhancers themselves. At the TE class level, LTR elements associated with cCREs are more likely to be distant from non-TE cCREs (Fig. 4A), which implies that LTRs are less dependent on genomic context in displaying regulatory activity compared to other TE classes. Lastly, we found that the distances from TEs associated with CTCF-only sites to non-TE CTCF-only sites are more consistent with random distribution in both human and mouse (Fig. 4A), despite abundant B2-derived CTCF binding sites in the mouse genome^25^. This indicates that CTCF binding sites provided by TEs are scattered randomly in the genome, which could facilitate formation of new chromatin boundaries. We performed the same analysis using mouse cCREs and TEs and found all trends observed in human to be consistent in the mouse genome (Extended Data Fig. 6A).

**Fig. 4.**
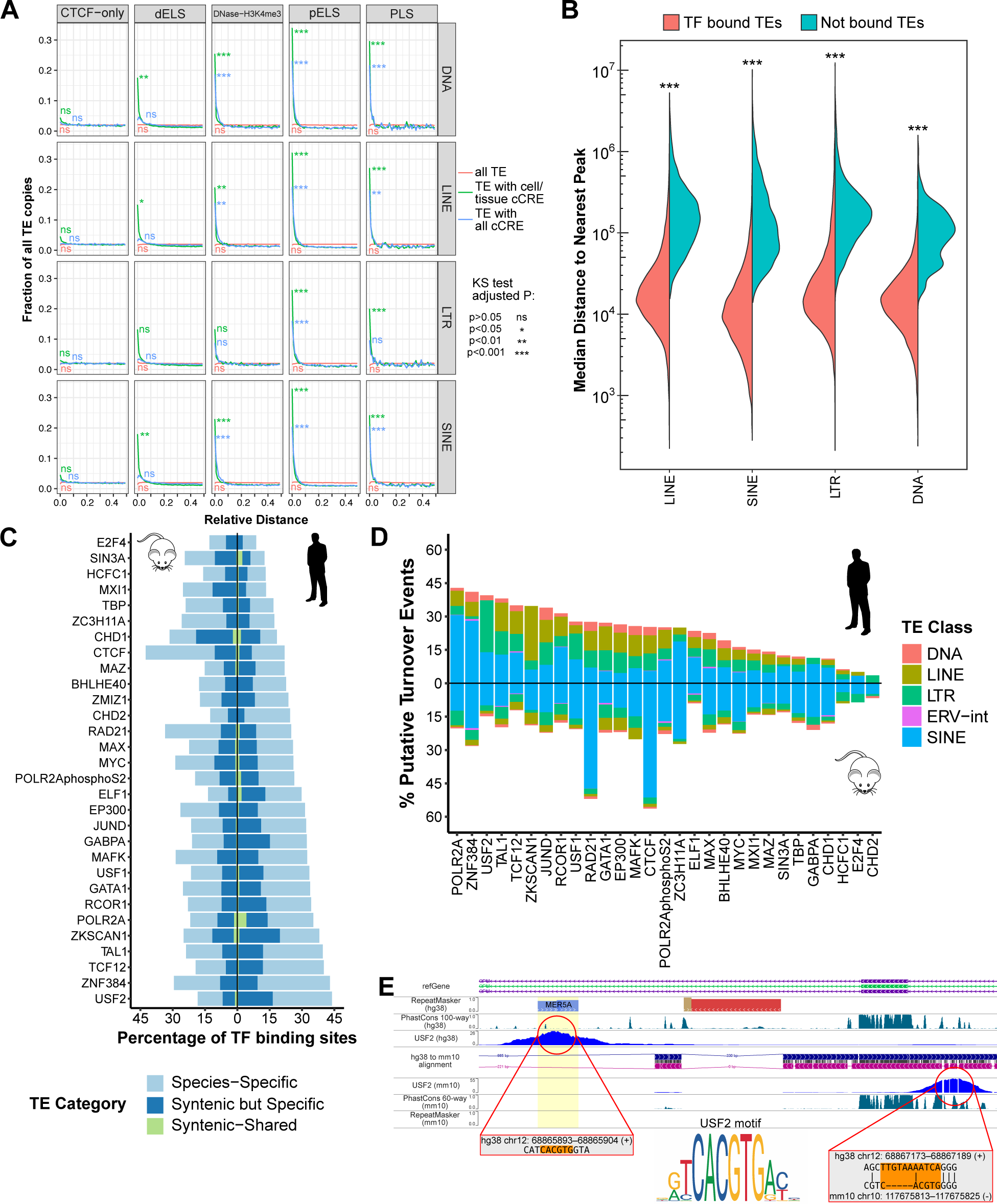
: Regulatory TEs cluster with non-TE regulatory elements and TEs provide TFBS turnover sites. A) Relative distance of all TEs to human cell agnostic cCREs (red), cCRE associated TEs to cell agnostic cCREs (blue), and cCRE associated TEs to same cell/tissue type cCREs (green). B) Median distances for TF bound TEs and non-bound TEs across 535 TF ChIP-seq datasets. C) Percentage of TE-derived TFBSs for 30 TFs with ChIP-seq in human K562 and mouse MEL cells. TE percentage is further divided into binding sites that are species-specific with no synteny, binding sites that are species-specific with synteny, and binding sites that are shared. D) Percentage of putative TFBS turnover events that come from TEs. Each percentage is split up by TE class contribution for the TF. E) Browser shot of USF2 binding site turnover in human facilitated by primate lineage insertion of MER5A. Underlying USF2 motif sequence alignment in human and mouse are shown (if available). *** p < 0.001

To further probe the connection between linear genomic distance and TE-derived *cis*-regulatory activity, we next examined the distance of TEs to TFBSs in K562 cells. For each of 535 TFs with ChIP-seq datasets where at least one TE subfamily was bound at least 10 times, we compared the distance of TF-bound TEs to their nearest non-TE TFBS of the same TF to that of non-bound TEs of the same subfamily. Across all TFs, we found that TF-bound TE copies are ∼10 times closer to non-TE TFBSs of the same TF than non-bound TE copies, regardless of TE class (Fig. 4B). These results are consistent with our distance analysis with cCREs and suggest that TEs with *cis*-regulatory activity tend to be proximal to other *cis*-regulatory elements.

Since TF-bound TEs tend to reside near non-TE TFBSs, we hypothesized that TEs can introduce local redundancy in TF binding, which may promote TFBS turnover during evolution^39–42^, whereby the TE-derived TFBS can functionally replace the nearby ancestral TFBS. To test this hypothesis, we selected 30 TFs with high quality ChIP-seq data in both human K562 and mouse MEL erythroleukemic cells, which are biologically homologous. As reported previously^43^, we found that up to ∼40% of TFBSs are contributed by TEs in each cell line (Fig. 4C). To identify putative TFBS turnover events, we searched for lineage-specific TFBSs within 5kb of a syntenic TFBS in the other lineage and inferred which TFBS was ancestral based on synteny and phastCons score (Extended Data Fig. 7A). Using this approach, we discovered a total of 6700 and 9245 putative TFBS turnover events across 30 TFs in human and mouse, respectively (Extended Data Fig. 7B). TEs make up 3-56% of putative turnover events, with most derived from lineage-specific TE insertions (Fig. 4D, 4E). The TFs with the highest TE-derived turnover rates are CTCF and RAD21 in mouse, both of which are part of the cohesin loading complex. These results are consistent with previous studies that have found TEs to participate in CTCF binding site turnover after human-mouse divergence^25^. Our findings point to TEs as important drivers of TFBS turnover during evolution.

## TE- and non-TE-derived cCREs have similar features

Since TEs contribute a large proportion of cCREs across the human genome, we explored whether TE-derived cCREs have distinct properties from non-TE cCREs. First, we considered sequence intrinsic *cis*-regulatory activity as measured by MPRA. Using ENCODE phase 4 lentivirus-based MPRA (lentiMPRA) data in K562 cells, which assayed all open chromatin sites in K562, we asked if the tested genomic sequences display differential regulatory activity based on TE annotation^44^. We classified sequences as TE-associated if at least 50% of the sequence is contributed by a single TE, resulting in ∼45,000 TE-associated and ∼76,000 non-TE sequences tested by MPRA. We further split sequences based on cCRE type. Overall, TE-associated sequences have similar or higher levels of MPRA activity compared to non-TE sequences of the same cCRE type (Fig. 5A). MPRA activity was significantly higher for TE-associated sequences in all cCRE types except PLS and CTCF-bound cCREs (Wilcoxon rank-sum test, pval < 0.005), but the differences in activity were subtle (median log_2_ fold change difference of 0.281, 0.048, 0.066, 0.082, and 0.041 for DNase-H3K4me3, dELS, pELS, DNase-only, and None, respectively). This is consistent with a previous study that found TE sequences of one LTR subfamily with higher MPRA activity than positive control sequences, albeit with far fewer tested elements^15^. Overall, these results suggest that TE-derived sequences possess the sequence potential for *cis*-regulatory activity as strong or stronger than their non-TE counterparts.

**Fig. 5:**
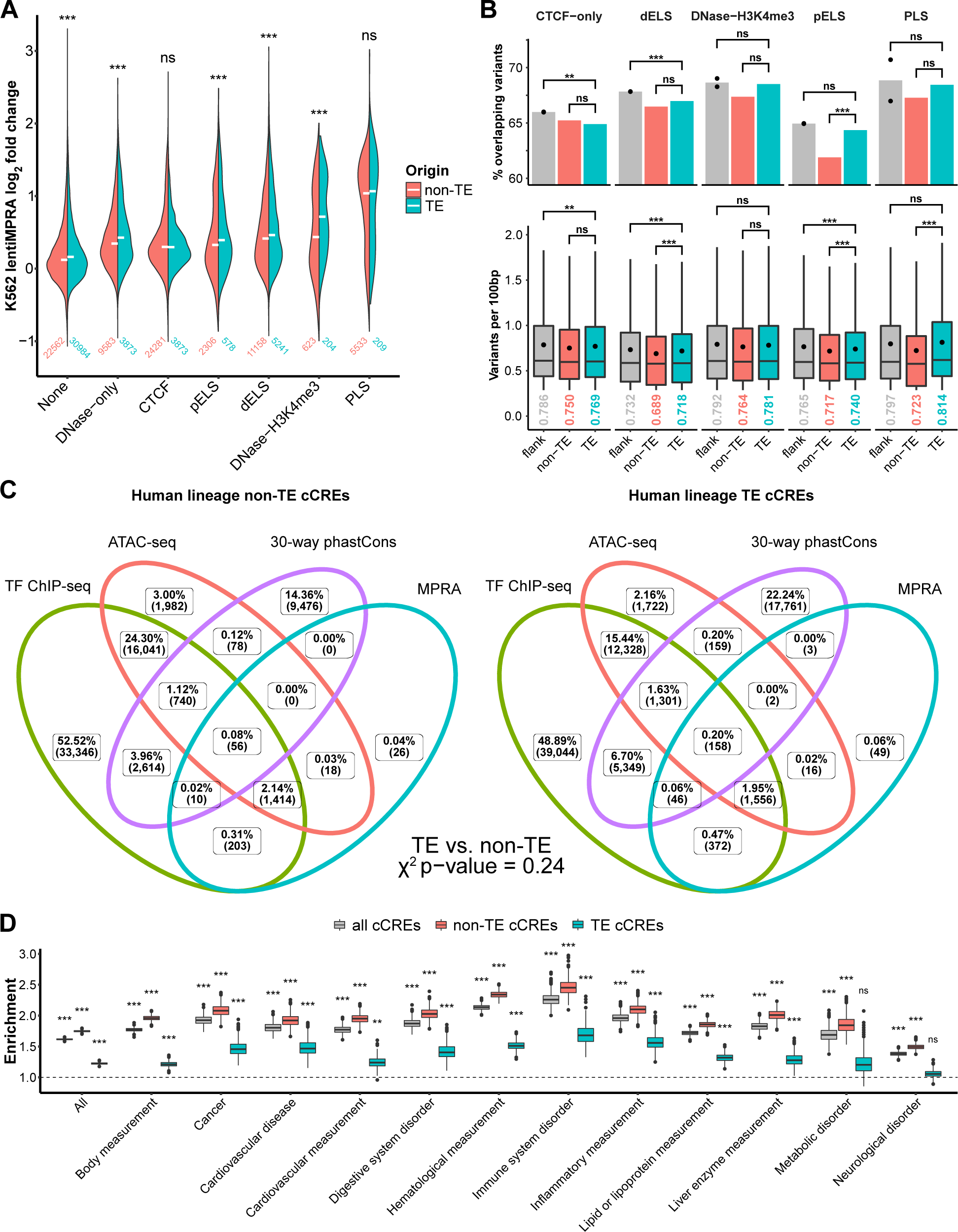
TE-derived cCREs share similar features with non-TE cCREs. A) K562 lentiMPRA activity split by cCRE annotation or lack thereof. Number of TE and non-TE sequences tested for each group is listed. Median log_2_ fold change over negative control activity is displayed as a white dash. Two-sided Wilcoxon rank sum test was performed to compare TE and non-TE sequence MPRA activity. B) 1000 Genomes Project common variants (allele frequency >1%) in TE and non-TE cCREs. Percentage of cCREs that overlap common variants (top) and variants per 100bp (bottom) are shown for each cCRE type. Percentages of common variant overlap for upstream and downstream cCRE flanking regions are shown as black dots (top). Mean variants per 100bp is displayed by a black dot and listed below each boxplot (bottom). Outliers have been removed from boxplots. Comparisons between TE cCREs, non-TE cCREs, and flanking regions were done using permutation tests. C) Venn diagram of human-lineage TE and non-TE cCRE overlap with TF ChIP-seq, ATAC-seq, 30-way phastCons, and MPRA activity. D) Enrichment of GWAS variants by EBI EFO parent term for all cCREs, non-TE cCREs, and TE cCREs. Permutation test was performed to compare observed overlap of cCREs with GWAS variants to shuffled genomic background. ** p < 0.01, *** p < 0.001, not significant (ns)

Next, we examined the frequency of variants found in the human population for TE-derived and non-TE cCREs based on the 1000 Genomes Project^45^. The expectation is that regions under functional constraint, like DNA elements regulating genes, would have fewer common variants, defined here as variants with human population allele frequency greater than 1%. Promoter distal TE-derived cCREs (dELS and CTCF-only) overlap variants less often than regions directly flanking them (Fig. 5B top, Extended Data Fig. 8). Furthermore, the frequency of common variants found in TE-derived cCREs is lower than their flanking regions apart from promoter sequences (PLS and DNase-H3K4me3) (Fig. 5B bottom, Extended Data Fig. 8). These results suggest that non-promoter TE-derived cCREs are under functional constraint and less tolerant of sequence variation than their surrounding sequences. However, TE-derived PLS, pELS, and dELS cCREs have significantly more common variants compared to non-TE cCREs, though the trend exists for all cCRE types. This indicates that TE-derived cCREs are generally less constrained in the human population than those apparently not derived from TEs.

Besides the epigenomic marks used by ENCODE to define cCREs, we used four additional metrics for identifying regulatory elements to compare the global profiles of TE-derived and non-TE cCREs: ATAC-seq for open chromatin, TF ChIP-seq for TF binding, MPRA for regulatory potential of the underlying sequence, and phastCons score for sequence conservation. As the vast majority of TE-derived cCREs are lineage-specific, we limited our analysis to TE-derived and non-TE cCREs that are found in human but not in mouse, allowing us to compare cCREs of roughly similar ages. Overall, TE-derived cCREs are not significantly different from non-TE cCREs in the proportion of elements that have any combination of ATAC-seq peaks, TF ChIP-seq peaks, MPRA activity, and high phastCons scores (Chi-square test, p = 0.24, Fig. 5C). This shows that the genomic features that are commonly used to annotate regulatory elements genome-wide are largely the same between TE and non-TE elements.

Finally, we investigated whether TE-derived cCREs could be physiologically relevant using the NHGRI-EBI GWAS catalog^46^. In addition to the comprehensive list of top GWAS SNPs (GWAS variants), we divided SNPs into different parent terms as defined by EBI for physiologically related diseases and traits. Compared to randomly shuffled genomic coordinates, the general set of cCREs is enriched for GWAS variants across all GWAS parent terms, in line with a previous study (Fig. 5D, Extended Data Fig. 9)^47^. TE-derived cCREs are enriched for GWAS variants overall and in 11/17 parent terms (Extended Data Fig. 9, Supplementary Table 2). However, they have consistently lower enrichment for GWAS variants compared to non-TE cCREs, which may be due to underrepresented profiling of SNPs in TEs from genotyping arrays (Supplementary Table 2). Altogether, these results suggest that TE-derived cCREs are functionally comparable to non-TE cCREs and carry sequences that are physiologically important for human traits and disease.

## Discussion

TEs make up a large portion of most mammalian genomes, and many studies have shown that TEs contribute to their regulatory landscape. However, the extent to which TEs supply different types of regulatory elements and the factors that allow them to evolve as regulatory elements are not well understood. In this study, we utilize cCREs to define the contribution of TEs to the human regulatory space, finding that ∼25% of all cCREs are TE-derived. This overall contribution is similar to previous estimates by Pehrsson et al. who studied the overlap of TEs with active regulatory states in the RoadMap Epigenome Project^22^. We observed that TEs do not contribute to the different types of cCREs equally; they contribute more substantially to gene-distal enhancers than to gene-proximal enhancers and promoters. This pattern is likely driven by selection against TE insertions in the proximity of genes^48^. Regardless of their cCRE type, we found that TE-derived cCREs are more likely to be restricted to one or a few cell/tissue types compared to non-TE cCREs. This result suggests that TEs could be important for regulatory innovation by providing gene regulatory elements that are active in a limited number of cellular contexts. The documented contribution of TEs to gene regulation in rapidly-evolving processes such as innate immunity and placentation support this hypothesis^13,49^.

Different TEs have invaded and expanded in genomes at various points during evolution, leading to some being shared between species and others being lineage-specific. We explored how TEs contribute to conserved and lineage-specific regulatory elements using cCREs annotated in human and mouse. While TEs provide a small fraction (up to 2%) of conserved cCREs orthologous between human and mouse, the vast majority of TE-derived cCREs are lineage-specific and account for 8-36% of all lineage-specific cCREs. Since cCREs have been less extensively profiled in mouse compared to human, it is possible that we are underestimating the contribution of TEs to conserved regulatory elements. We next showed that most TEs that are orthologous (shared by descent) between human and mouse are retained as or become cCREs in only one lineage, indicating either lineage-specific loss or lineage-specific gain of regulatory activity, respectively. In addition to most non-orthologous, lineage-specific TE-cCREs coming from TE subfamilies that are old enough to be found in both human and mouse (Extended Data Fig. 2), our results suggest that most TE-derived regulatory elements come from old TEs. This is consistent with a previous study by Villar et al. which found that evolutionarily young enhancers were primarily adapted from ancestral DNA sequences over 100 million years of age^50^.

To broadly understand where *cis*-regulatory activity in TEs comes from, we investigated the evolutionary origins of cCRE-associated TF motifs. In LINEs, LTR elements, and DNA transposons, cCRE-associated TF motifs originate from their consensus sequences more than expected by chance, suggesting an ancestral source for many important TE-derived TF motifs. As TEs age, the bulk of these TF motifs degrade over time. These results suggest that the amplification of these TEs disperses TF motifs available immediately upon insertion, but only a small subset is co-opted for host gene regulation. SINEs, which are extremely abundant in mammals (Alu elements alone account for 10% of the human genome sequence), show a very different trend compared to the other main TE classes. They have the lowest proportion of cCRE-associated TF motifs stemming from their consensus sequence, and the percentage of motifs that do not have an ancestral origin does not decrease as they age. These results suggest that SINEs provide relatively fewer mature TF motifs but instead frequently supply raw sequence material from which additional TF motifs emerge over time by mutation. This model is consistent with previous studies documenting the progressive birth of enhancers from Alu and RSINE1 elements in human and mouse, respectively^18,51^. Since SINEs have given rise to ∼5% of human cCREs, these findings indicate that this “seed-and-mature” process has been a rich source of new *cis*-regulatory elements in the human genome.

When TEs insert themselves into the genome, the newly integrated copy and its progenitor are typically identical in sequence and therefore have the same sequence potential for *cis*-regulatory activity. However, only a small subset of TE copies within any given subfamily overlaps with cCREs. What influences some TE copies to retain or evolve *cis*-regulatory activity? One likely influential factor is the genomic context of the TE and its proximity to functional sequences such as genes and *cis*-regulatory elements^18^. Consistent with this model, we demonstrate that TEs with either cCREs or TFBSs have shorter genomic distance to non-TE cCREs or TFBSs compared to other TEs. Based on this observation, we propose that the clustering of TFBSs from TE insertions introduces functional redundancy that can promote the turnover of TFBSs during evolution, as is prominent in mammals^39–42^. By examining the binding profiles of 30 TFs in human and mouse leukemia cell lines, we estimate that TEs have contributed 3-56% of all putative TFBS turnover events depending on the TF. Together these results suggest that the insertion of TEs near existing *cis*-regulatory elements is a major driver of TFBS evolutionary turnover.

An outstanding question for future studies is to determine the extent to which TE-derived cCREs have contributed to human adaptation and phenotypic variation. Our analysis brings hints that a subset of TE-derived cCREs serve important biological functions. First, we found that the sequences of TE-derived cCREs are generally more evolutionarily constrained than their non-cCRE counterparts. Second, we observed that TE-derived cCREs are enriched for GWAS variants, albeit to a lesser extent than non-TE cCREs. While this could indicate that TEs are less likely to be physiologically relevant, it could also reflect technical shortcomings associated with genotyping within TE sequences. Genotyping arrays, which use short oligonucleotide probes to discern SNPs, are designed to avoid repetitive regions of the genome. Our analysis of nine genotyping arrays from Affymetrix and Illumina shows between 30-36% of SNPs located in repetitive DNA, short of the 45% TE content in the human genome (Supplementary Table 2). These observations suggest that GWAS studies may have missed trait-associated SNPs residing within TE sequences and there is a need to consider TE-derived variants in follow-up studies to GWAS^52^. Our study confirms that TEs are responsible for a substantial level of *cis*-regulatory activity in the human genome and thus hold important keys to our understanding of human biology and disease.

## Methods

### Annotation of TE-derived cCREs

Genomic cCRE annotations in hg38 (cell agnostic and 25 fully profiled ENCODE cell/tissue types) and mm10 were downloaded from (https://screen.wenglab.org/) and the ENCODE portal (https://www.encodeproject.org/)^19^. Genomic TE annotations in hg38 and mm10 were obtained from RepeatMasker (https://repeatmasker.org/)^53^. We used BedTools intersect^54^ to find cCREs that are associated with TEs, requiring at least 50% of the cCRE to overlap a single TE.

### Enrichment of TEs in cCREs

We calculated the enrichment of TE subfamilies for cCREs as follows.

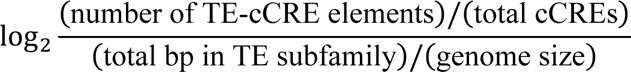

For visualization, we included TE subfamilies with no overlap with cCREs as log_2_ enrichment of −10, which is lower than any subfamily with cCRE overlap.

### Human-mouse cCRE comparison

To characterize human and mouse cCREs as shared or lineage-specific (Extended Data Fig. 2), we first used liftOver with -minMatch option of 0.1 to identify syntenic regions in the other species. The syntenic region was determined to be a cCRE or TE-derived if at least 50% of the syntenic region overlaps with a cCRE or TE. Syntenic regions with cCREs were classified as “shared” if the cCRE type was the same in both species and “different” if the cCRE type was different. TEs in syntenic regions of human and mouse were counted as orthologous if both TEs are annotated as belonging to the same TE family (e.g. SINE/Alu).

To compare sequence conservation, 100-way phastCons scores^55^ were downloaded from https://genome.ucsc.edu/. Two-sided Wilcoxon rank sum test was used for comparisons between groups of TE-cCREs.

### Identification of cCRE-enriched TF motifs

First, TEs in each subfamily were separated based on overlap with hg38 cCREs, with subfamilies that lacked cCRE overlap removed from analysis (n=116). Next, TE subfamilies were split into three groups depending on whether the length distributions of cCRE (foreground) and non-cCRE (background) elements were significantly different based on Kolmogorov-Smirnov (KS) test. One group of subfamilies (n=194) have no significant difference with all background elements included. For the second group of subfamilies (n=993) with significant difference in length distribution between foreground and background elements, background elements were binned and randomly selected to match the proportion of foreground elements found in each bin. Random selection of background elements in the second group of subfamilies was performed 10 times. TE subfamilies that could not achieve matched foreground/background length distributions were disregarded from further analysis (n=17).

To identify cCRE-enriched motifs, we ran AME motif enrichment using the HOCOMOCOv11 human core transcription factor motif database^56^ for each TE subfamily, with cCRE elements as foreground and non-cCRE elements (all elements or random selection) as background/control. Enriched motifs were grouped according to motif archetypes^47^. To confirm AME results, we scanned TE subfamily elements for the top enriched motif within each archetype and performed Fisher’s exact test, further filtering for motifs that have significant association with cCRE annotation (p < 0.05 after multiple test correction with Benjamini-Hochberg method), at least 10 elements having both the motif and cCRE annotation, and odds ratios of at least 2. We also filtered for TF motifs that pass Fisher’s exact test using TE subfamily consensus coverage-controlled background sequences, identifying TF motifs that distinguish cCRE overlapping TE copies from non-overlapping copies based on sequence variation alone.

### Origin of cCRE-associated TF motifs

In order to estimate the percentage of cCRE enriched motifs that were derived from an ancestral origin, we first derived consensus sequences for each TE subfamily from RepeatMasker and the RepBase-derived RepeatMasker Library 20170127 (Supplementary Methods). We could not obtain consensus sequences for four subfamilies (L2d, L2d2, Alu, and MLT1B-int), which were excluded from further analysis. Next, we scanned each consensus sequence for all HOCOMOCOv11 human core motifs. For each motif found in a TE subfamily’s consensus sequence, we scanned all elements within the subfamily for the motif. The relative position of each motif to the consensus sequence was found by aligning each element to its consensus sequence using Needle pairwise alignment^57^. Finally, the percent ancestral origin rate of a given motif was calculated as the percentage of motifs that were within 10bp of the consensus sequence motif. As we had grouped motifs based on motif archetype, we used the ancestral origin rate of the top enriched motif per archetype. In the case that the top motif was not found in the consensus sequence, we allowed for any other enriched motif in the archetype that was in the consensus to substitute. Any motif archetype that had no cCRE enriched motif in the consensus sequence was assigned an ancestral origin rate of 0.

### Relative distance of TE to closest cCRE

To estimate the spatial correlation between TEs and cCREs, we calculated relative distance first described by Favorov et al. and implemented within the BEDTools suite^54,58^. Briefly, for each cCRE type, TEs were assigned to their closest non-TE cCRE. Then, the distance between cCREs was split into 100 equal sized intervals and the frequency of TEs that fall within each interval was counted. We shuffled TE coordinates using bedtools shuffle 100 times to constitute the null hypothesis set, and applied KS test with Bonferroni multiple test correction to evaluate the difference between observation and shuffled null expectation.

### Bound vs unbound TE distance to nearest TF peak

A total of 587 IDR thresholded TF ChIP-seq peak files in K562 were downloaded from ENCODE after filtering out those with “NOT_COMPLIANT” or “ERROR” audit labels. For each TF, individual TEs were classified as bound if they intersected the peak summit and non-bound otherwise. We then calculated the linear distance from each TE to the nearest non-overlapping peak. For each TE subfamily with at least 10 TF bound individual TEs, we randomly sampled an equal number of non-bound individual TEs as those which were bound and ranked them in descending order of distance. After repeating random sampling 1000 times, we averaged each of the ranks across all 1000 samples to get a distribution of average distances to the nearest non-overlapping peak for non-bound TEs. We then calculated p-values using two-sided Wilcoxon rank sum tests between bound and unbound TEs within each subfamily. We calculated the log_10_ ratio of average median distances to the nearest non-overlapping peak between bound and non-bound TEs as the following:

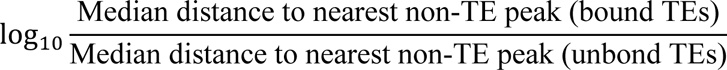

### Identification of putative TF turnover events

IDR-thresholded peaks for 30 TF ChIP-seq datasets with matching K562 TF ChIP-seq were downloaded for mouse MEL from ENCODE^19^. Syntenic regions to TF binding peaks were identified with the same method described above for human-mouse comparison. If a TF peak in one species overlapped at least 50% of a peak in the other species, it was classified as “shared”. Otherwise, the TF peak was classified as “syntenic but specific” for alignable sequences but with species specific TF binding. To identify putative TFBS turnover events after human-mouse divergence, we identified all TF peaks in one species within 5kb of the syntenic region of a TF peak in the other species (Extended Data Fig. 7A). For each peak, mean phastCons score was assigned using 100-way vertebrate phastCons scores in human or 60-way vertebrate phastCons scores in mouse^55^. We calculated the median phastCons score for conserved TF binding peaks in human and mouse to infer human-mouse ancestral TF binding. For each pair of lineage-specific peaks, the human-mouse ancestral TFBS was inferred based on human-mouse synteny and phastCons score higher than the median phastCons score for conserved TF binding peaks. Pairs of lineage-specific peaks were identified as putative TFBS turnover events if a single non-ancestral TF binding peak was within 5kb of an ancestral TF binding peak. TE-derived peaks were classified using the prior mentioned criteria of 50% overlap with the TF binding peak.

### MPRA comparison

K562 lentiMPRA data was downloaded from the ENCODE portal^44^. TE-derived or non-TE-derived cCRE annotations were intersected with lentiMPRA sequence coordinates and then assigned the maximum log_2_ fold change (log2FC) value (both strands). cCREs were grouped based on their annotation with the following exceptions: “None” = Low-DNase or no intersection, “CTCF” = any classification bound by CTCF. Two-sided Wilcoxon rank sum test was performed comparing the log2FC values of TE-derived cCREs with non-TE-derived cCREs within the same category. An alternative hypothesis of TE-derived cCREs having a greater log2FC value than non-TE-derived cCREs was used. P-values for all tests underwent Benjamini-Hochberg multiple-test correction.

### Human population variant frequency in cCREs

We extracted variants within the human population characterized by the 1000 genomes project^45^ and further selected variants with allele frequency >1% as common variants. For each cCRE that did not overlap coding sequence in GENCODEv41^59^, we counted how many common variants intersect with them, and the number of common variants was normalized per 100bp of sequence. The percentage with variant overlap and the distribution of variants per 100bp was obtained for TE-derived cCREs, non-TE cCREs, and cCRE flanking regions. Flanking regions were defined as non-coding genomic regions directly upstream and downstream with the same length as the cCRE. Permutation tests were then performed to compare percentage with variant overlap and mean variants per 100bp between TE-derived cCREs, non-TE cCREs, and cCRE flanking regions relative to random genomic background (Supplementary Methods).

### Venn Diagram comparing features for TE-derived and non-TE-derived cCREs

ATAC-seq in K562 cells were downloaded from ENCODE and 30-way (27 primates) phastCons scores were downloaded from https://genome.ucsc.edu/. Non-TE and TE-derived cCREs were classified as being accessible (ATAC-seq), bound by a TF (TF ChIP-seq), MPRA active (MPRA), or having high levels of sequence conservation among primates (phastCons). PhastCons scores were binned into 20bp windows and each cCRE was assigned the mean of intersecting phastCons scores. cCREs with the top 10% of phastCons scores were selected as high sequence conservation. For ATAC-seq and TF ChIP-seq, a cCRE containing a peak summit within its interval was considered accessible or bound, respectively. MPRA log2FC was obtained for each cCRE as previously described, and cCREs were classified as active in MPRA if the maximum log2FC was greater than 1. Finally, a Venn Diagram was generated using the combined classification of cCREs. Chi-square test was performed to test for differences in feature classification between TE- and non-TE-derived cCREs.

### Enrichment of cCREs in GWAS variants

The NHGRI-EBI GWAS catalog with added ontology annotations and GWAS to EFO mappings was downloaded from http://www.ebi.ac.uk/gwas^46^. The strongest SNP was chosen for each reported entry. We used dbSNP153^60^ to assign chromosome positions in hg38 to SNPs if chromosome position was not already listed; SNPs with neither chromosome position nor dbSNP153 rs ID were excluded. The number of GWAS SNPs found in cCREs was counted following BEDTools overlap^54^. Each SNP was counted at most once for each parent term. Permutation test by genome-wide shuffling of cCRE coordinates was performed 1,000 times to obtain a random expectation for GWAS SNP overlap. Enrichment was calculated as the number of overlapping GWAS variants in cCREs divided by the number of overlapping GWAS variants in random shuffled coordinates. P-value was defined as the proportion of random shuffles that reached the number of overlapping GWAS variants in cCREs or higher. As a negative control, we shuffled cCRE coordinates an additional 100 times and took the mean number of GWAS variant overlaps.

## Data availability

All accession codes and download links for publicly available data are listed in Supplementary Table 1.

## Code availability

All code for analysis is available upon request.

## Acknowledgments

This work was supported by NIH grants R01HG007175 and U01HG009391. A.Y.D. was supported by NHGRI training grant T32HG000045. J.D.C. was supported by NIGMS MIRA (2R35GM122550-06).

## Author contributions

T.W. and C.F. conceived the project. A.Y.D., J.D.C., and X.Z. designed and performed analysis.

All authors took part in writing the manuscript.

## Corresponding authors

Correspondence to Ting Wang or Cédric Feschotte.

## References

1. McClintock, B. The Origin and Behavior of Mutable Loci in Maize. Proc. Natl. Acad. Sci. U. S. A. 36, 344 (1950).

2. Osmanski, A. B. et al. Insights into mammalian TE diversity through the curation of 248 genome assemblies. Science (80-.). 380, (2023).

3. Nurk, S. et al. The complete sequence of a human genome. Science (80-.). 376, 44–53 (2022).

4. Wells, J. N. & Feschotte, C. A Field Guide to Eukaryotic Transposable Elements. Annu. Rev. Genet. 54, 539–561 (2020).

5. Christmas, M. J. et al. Evolutionary constraint and innovation across hundreds of placental mammals. Science (80-.). 380, (2023).

6. Rebollo, R., Romanish, M. T. & Mager, D. L. Transposable Elements: An Abundant and Natural Source of Regulatory Sequences for Host Genes. Annu. Rev. Genet. 46, 21–42 (2012).

7. Chuong, E. B., Elde, N. C. & Feschotte, C. Regulatory activities of transposable elements: from conflicts to benefits. Nat. Rev. Genet. 18, 71–86 (2017).

8. Bourque, G. et al. Ten things you should know about transposable elements. Genome Biol. 19, 199 (2018).

9. Sundaram, V. & Wysocka, J. Transposable elements as a potent source of diverse cis-regulatory sequences in mammalian genomes. Philos. Trans. R. Soc. B 375, (2020).

10. Fueyo, R., Judd, J., Feschotte, C. & Wysocka, J. Roles of transposable elements in the regulation of mammalian transcription. Nat. Rev. Mol. Cell Biol. 2022 237 23, 481–497 (2022).

11. Lawson, H. A., Liang, Y. & Wang, T. Transposable elements in mammalian chromatin organization. Nat. Rev. Genet. 2023 1–12 (2023). doi:10.1038/s41576-023-00609-6

12. Wang, T. et al. Species-specific endogenous retroviruses shape the transcriptional network of the human tumor suppressor protein p53. Proc. Natl. Acad. Sci. U. S. A. 104, 18613–8 (2007).

13. Chuong, E. B., Elde, N. C. & Feschotte, C. Regulatory evolution of innate immunity through co-option of endogenous retroviruses. Science 351, 1083–7 (2016).

14. Sundaram, V. et al. Functional cis-regulatory modules encoded by mouse-specific endogenous retrovirus. Nat. Commun. 8, (2017).

15. Du, A. Y. et al. Functional characterization of enhancer activity during a long terminal repeat’s evolution. Genome Res. 32, 1840–1851 (2022).

16. Zemojtel, T., Kielbasa, S. M., Arndt, P. F., Chung, H. R. & Vingron, M. Methylation and deamination of CpGs generate p53-binding sites on a genomic scale. Trends Genet. 25, 63–66 (2009).

17. Zemojtel, T. et al. CpG Deamination Creates Transcription Factor–Binding Sites with High Efficiency. Genome Biol. Evol. 3, 1304–1311 (2011).

18. Judd, J., Sanderson, H. & Feschotte, C. Evolution of mouse circadian enhancers from transposable elements. Genome Biol. 2021 221 22, 1–26 (2021).

19. The ENCODE Project Consortium et al. Expanded encyclopaedias of DNA elements in the human and mouse genomes. Nature 583, 699–710 (2020).

20. Roadmap Epigenomics Consortium et al. Integrative analysis of 111 reference human epigenomes. Nat. 2015 5187539 518, 317–330 (2015).

21. Trizzino, M., Kapusta, A. & Brown, C. D. Transposable elements generate regulatory novelty in a tissue-specific fashion. BMC Genomics 19, (2018).

22. Pehrsson, E. C., Choudhary, M. N. K., Sundaram, V. & Wang, T. The epigenomic landscape of transposable elements across normal human development and anatomy. Nat. Commun. 2019 101 10, 1–16 (2019).

23. Brocks, D. et al. DNMT and HDAC inhibitors induce cryptic transcription start sites encoded in long terminal repeats. Nat. Genet. 49, 1052–1060 (2017).

24. Schmidt, D. et al. Waves of retrotransposon expansion remodel genome organization and CTCF binding in multiple mammalian lineages. Cell 148, 335–348 (2012).

25. Choudhary, M. N. K. et al. Co-opted transposons help perpetuate conserved higher-order chromosomal structures. Genome Biol. 21, 1–14 (2020).

26. Choudhary, M. N. K., Quaid, K., Xing, X., Schmidt, H. & Wang, T. Widespread contribution of transposable elements to the rewiring of mammalian 3D genomes. Nat. Commun. 2023 141 14, 1–12 (2023).

27. Simonti, C. N., Pavličev, M. & Capra, J. A. Transposable Element Exaptation into Regulatory Regions Is Rare, Influenced by Evolutionary Age, and Subject to Pleiotropic Constraints. Mol. Biol. Evol. 34, 2856 (2017).

28. Diehl, A. G., Ouyang, N. & Boyle, A. P. Transposable elements contribute to cell and species-specific chromatin looping and gene regulation in mammalian genomes. Nat. Commun. 2020 111 11, 1–18 (2020).

29. Kuhn, R. M., Haussler, D. & James Kent, W. The UCSC genome browser and associated tools. Brief. Bioinform. 14, 144–161 (2013).

30. Chinwalla, A. T. et al. Initial sequencing and comparative analysis of the mouse genome. Nature 420, 520–562 (2002).

31. Jordan, I. K., Rogozin, I. B., Glazko, G. V & Koonin, E. V. Origin of a substantial fraction of human regulatory sequences from transposable elements. Trends Genet. 19, 68–72 (2003).

32. Van De Lagemaat, L. N., Landry, J. R., Mager, D. L. & Medstrand, P. Transposable elements in mammals promote regulatory variation and diversification of genes with specialized functions. Trends Genet. 19, 530–536 (2003).

33. Lowe, C. B., Bejerano, G. & Haussler, D. Thousands of human mobile element fragments undergo strong purifying selection near developmental genes. Proc. Natl. Acad. Sci. U. S. A. 104, 8005–10 (2007).

34. Feschotte, C. Transposable elements and the evolution of regulatory networks. Nat. Rev. Genet. 9, 397–405 (2008).

35. Swergold, G. D. Identification, Characterization, and Cell Specificity of a Human LINE-1 Promoter. Mol. Cell. Biol. 10, 6718–6729 (1990).

36. Minakami, R. et al. Identification of an internal cis-element essential for the human Li transcription and a nuclear factor(s) binding to the element. Nucleic Acids Res. 20, 3139– 3145 (1992).

37. Alexandrova, E. A. et al. Sense transcripts originated from an internal part of the human retrotransposon LINE-1 5′ UTR. Gene 511, 46–53 (2012).

38. Sun, X. et al. Transcription factor profiling reveals molecular choreography and key regulators of human retrotransposon expression. Proc. Natl. Acad. Sci. U. S. A. 201722565 (2018). doi:10.1073/pnas.1722565115

39. Stefflova, K. et al. Cooperativity and Rapid Evolution of Cobound Transcription Factors in Closely Related Mammals. Cell 154, 530–540 (2013).

40. Yue, F. et al. A comparative encyclopedia of DNA elements in the mouse genome. Nat. 2014 5157527 515, 355–364 (2014).

41. Cheng, Y. et al. Principles of regulatory information conservation between mouse and human. Nat. 2014 5157527 515, 371–375 (2014).

42. Vierstra, J. et al. Mouse regulatory DNA landscapes reveal global principles of cis-regulatory evolution. Science (80-.). 346, 1007–1012 (2014).

43. Sundaram, V. et al. Widespread contribution of transposable elements to the innovation of gene regulatory networks. Genome Res. 24, 1963–76 (2014).

44. Agarwal, V. et al. Massively parallel characterization of transcriptional regulatory elements in three diverse human cell types. bioRxiv 2023.03.05.531189 (2023). doi:10.1101/2023.03.05.531189

45. The 1000 Genomes Project Consortium. A global reference for human genetic variation. Nature 526, 68–74 (2015).

46. Sollis, E. et al. The NHGRI-EBI GWAS Catalog: knowledgebase and deposition resource. Nucleic Acids Res. 51, D977–D985 (2023).

47. Vierstra, J. et al. Global reference mapping of human transcription factor footprints. Nat. 2020 5837818 583, 729–736 (2020).

48. Medstrand, P., Van De Lagemaat, L. N. & Mager, D. L. Retroelement Distributions in the Human Genome: Variations Associated With Age and Proximity to Genes. Genome Res. 12, 1483–1495 (2002).

49. Lynch, V. J., Leclerc, R. D., May, G. & Wagner, G. P. Transposon-mediated rewiring of gene regulatory networks contributed to the evolution of pregnancy in mammals. Nat Genet 43, 1154–1159 (2011).

50. Villar, D. et al. Enhancer evolution across 20 mammalian species. Cell 160, 554–566 (2015).

51. Su, M., Han, D., Boyd-Kirkup, J., Yu, X. & Han, J. D. J. Evolution of Alu Elements toward Enhancers. Cell Rep. 7, 376–385 (2014).

52. Payer, L. M. et al. Structural variants caused by Alu insertions are associated with risks for many human diseases. Proc. Natl. Acad. Sci. U. S. A. 114, E3984–E3992 (2017).

53. Smit, A., Hubley, R. & Green, P. RepeatMasker Open-4.0. 2013-2015 <http://www.repeatmasker.org>

54. Quinlan, A. R. & Hall, I. M. BEDTools: a flexible suite of utilities for comparing genomic features. Bioinformatics 26, 841–842 (2010).

55. Siepel, A. et al. Evolutionarily conserved elements in vertebrate, insect, worm, and yeast genomes. Genome Res. 15, 1034–1050 (2005).

56. Kulakovskiy, I. V. et al. HOCOMOCO: towards a complete collection of transcription factor binding models for human and mouse via large-scale ChIP-Seq analysis. Nucleic Acids Res. 46, D252–D259 (2018).

57. Needleman, S. B. & Wunsch, C. D. A general method applicable to the search for similarities in the amino acid sequence of two proteins. J. Mol. Biol. 48, 443–453 (1970).

58. Favorov, A. et al. Exploring Massive, Genome Scale Datasets with the GenometriCorr Package. PLOS Comput. Biol. 8, e1002529 (2012).

59. Frankish, A. et al. GENCODE 2021. Nucleic Acids Res. 49, D916–D923 (2021).

60. Sherry, S. T., Ward, M. & Sirotkin, K. dbSNP—Database for Single Nucleotide Polymorphisms and Other Classes of Minor Genetic Variation. Genome Res. 9, 677–679 (1999).

