## Extended Data Figures for "Regulatory Transposable Elements in the Encyclopedia of DNA Elements"

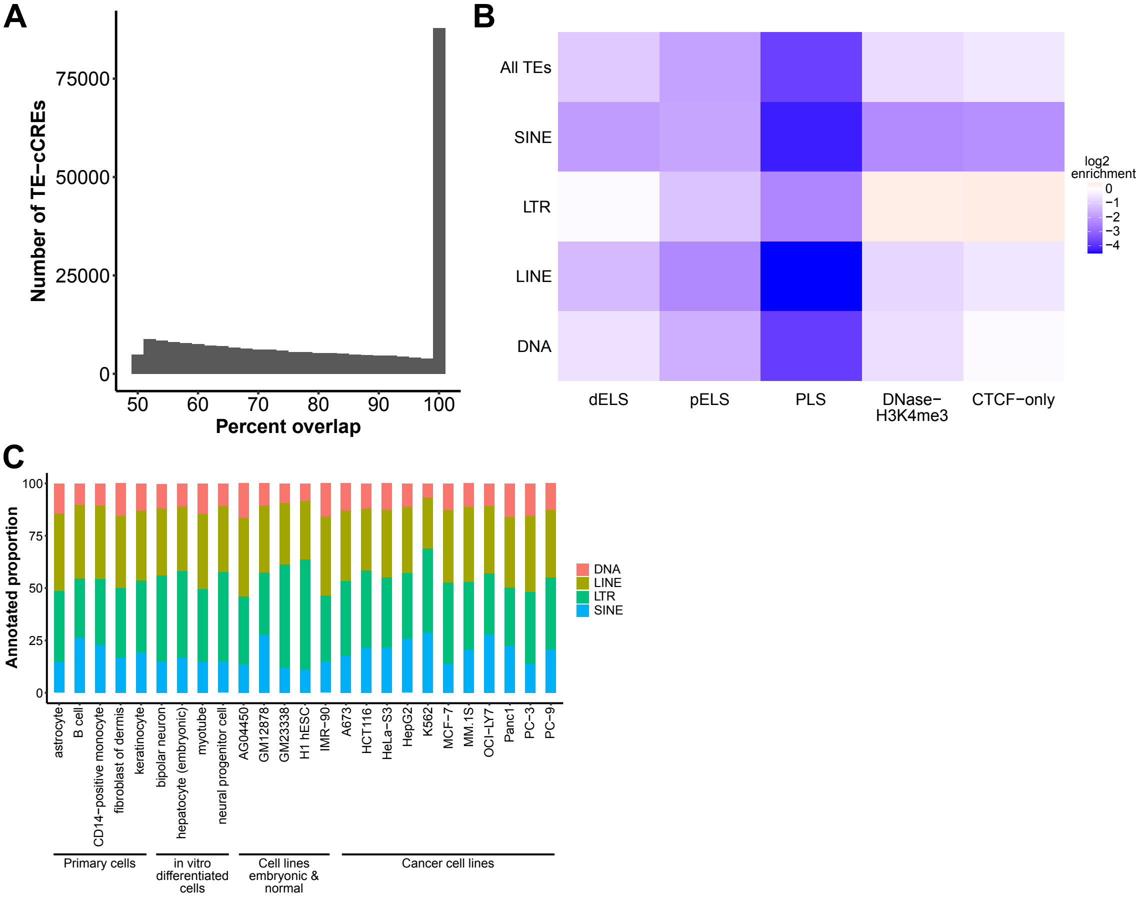


**Extended Data Fig. 1**: Overlap of TEs with human cCREs. A) Percentage of TE-derived cCREs that overlap a single TE. B) Log2 enrichment of TE class overlap with cCREs. C) Percentage of TE-derived cCREs from each TE class across 25 fully classified cell/tissue types.

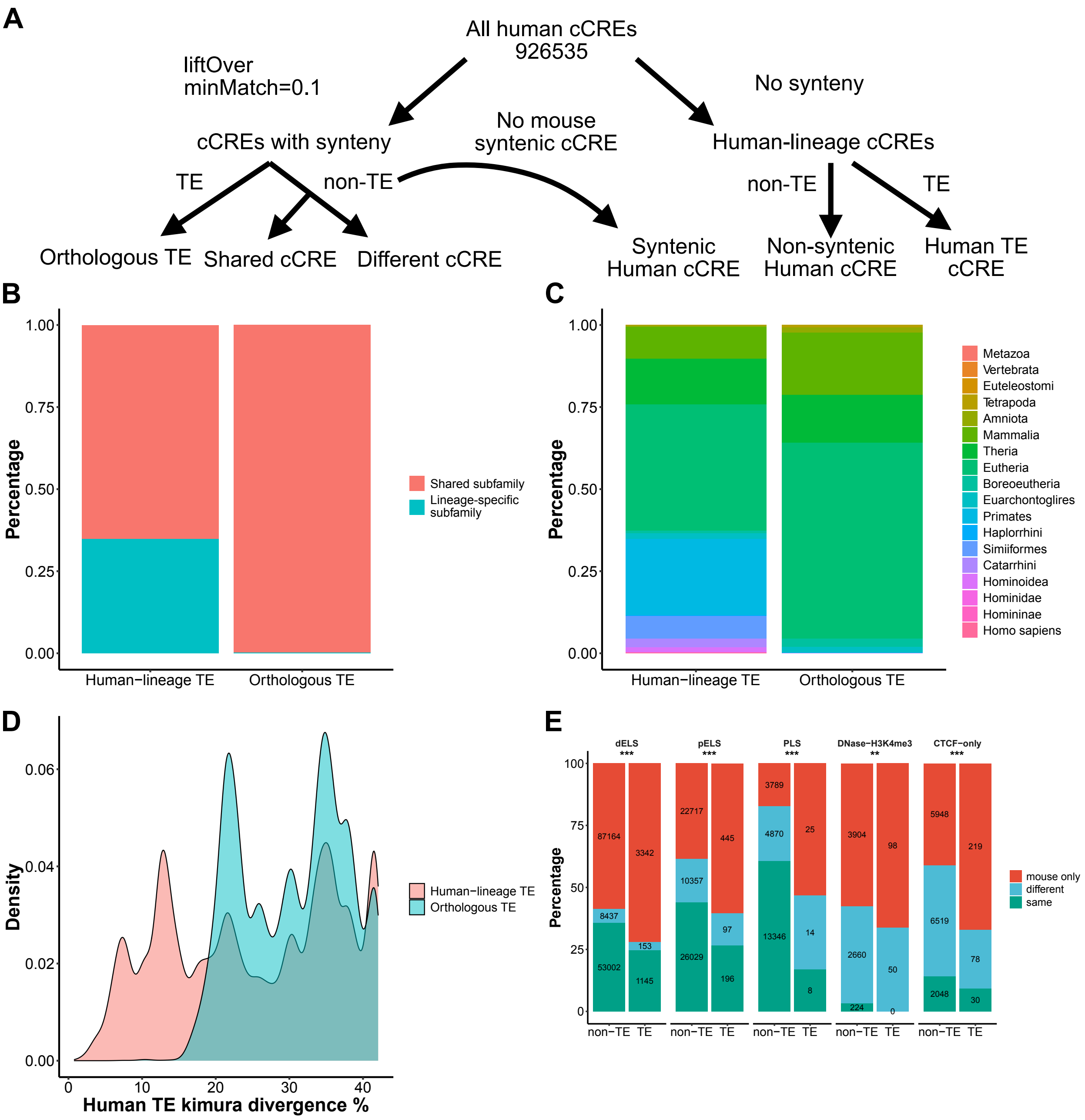


**Extended Data Fig. 2**: TE-derived conserved and lineage-specific cCREs in human and mouse. A) Schematic for defining orthologous TEs, conserved cCREs, and human-lineage cCREs. B) Percentage of orthologous TEs which overlap human cCREs and human-lineage TE-cCREs that come from human-mouse shared and lineage-specific TE subfamilies. C) Percentage of orthologous TEs which overlap human cCREs and human-lineage TE-cCREs based on clade. D)

TE subfamily Kimura divergence distribution for orthologous TEs which overlap human cCREs and human-lineage TE-cCREs. E) Percentage of cCREs that are shared or lineage-specific for orthologous TE and syntenic non-TE mouse anchored cCRE regions. Shared cCREs are split into "same" and "different" categories depending on the syntenic mouse and human cCRE types. Grouping by cCRE type is done using the anchored mouse cCRE. ** $p < 0.01$, *** $p < 0.001$

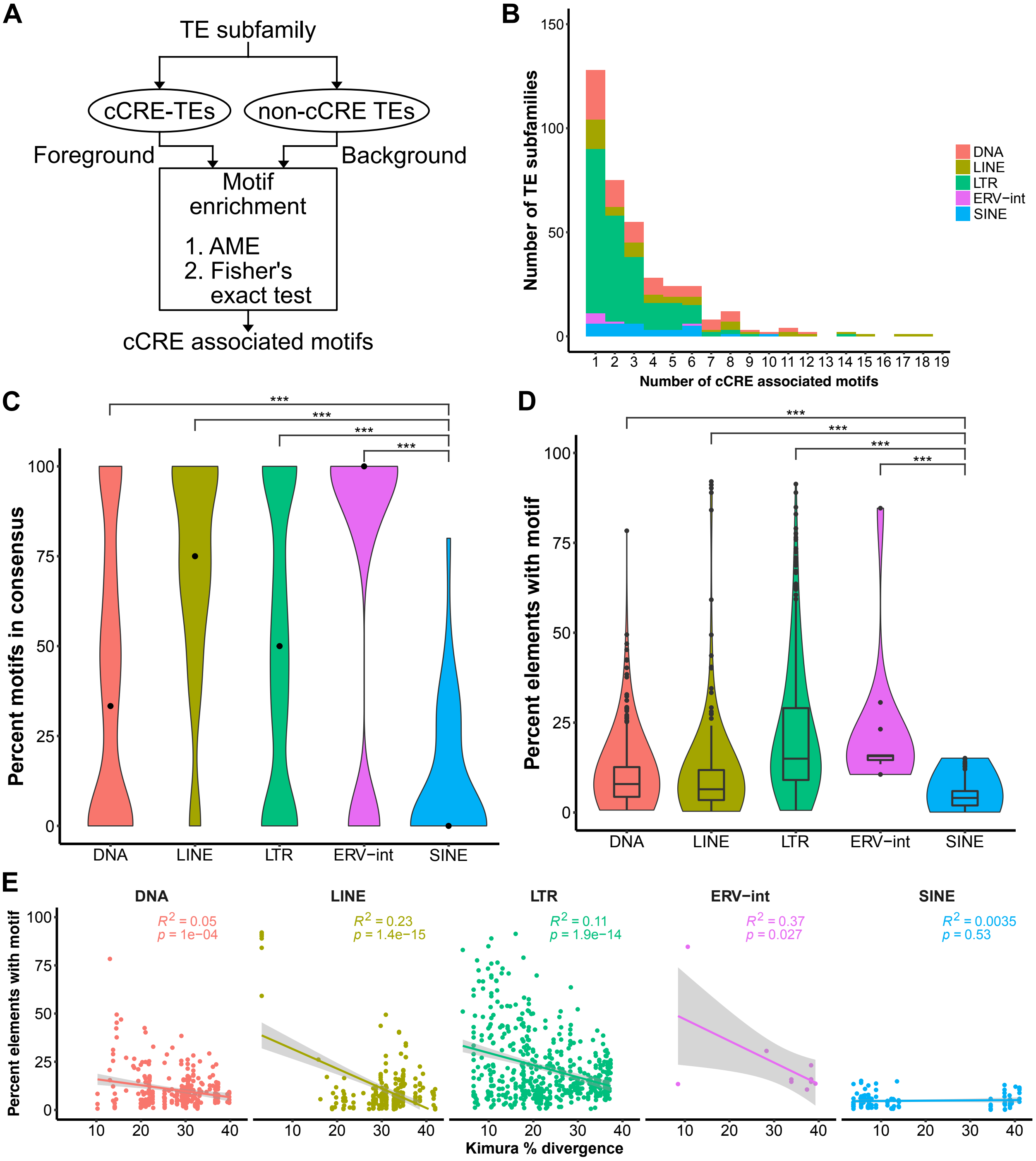


**Extended Data Fig. 3**: Origin of cCRE associated TF motifs in TEs. A) Schematic for identification of cCRE associated TF motifs per TE subfamily. B) Distribution of cCRE associated motif number for TE subfamilies with at least one associated motif. C) Percentage of cCRE associated motifs that are found in the TE subfamily consensus sequence, per TE subfamily and grouped by TE class. Median percentages are indicated by a black dot. Comparisons to the SINE class by Wilcoxon rank-sum test are shown. D) Percentage of TE subfamily elements that harbor a given cCRE associated motif. A subfamily with multiple motifs has one datapoint for each motif. Comparisons to the SINE class by Wilcoxon rank-sum test are shown. E) Relationship between percentage of TE subfamily elements that harbor a given cCRE associated motif and TE subfamily age approximated by Kimura divergence. *** $p < 0.001$

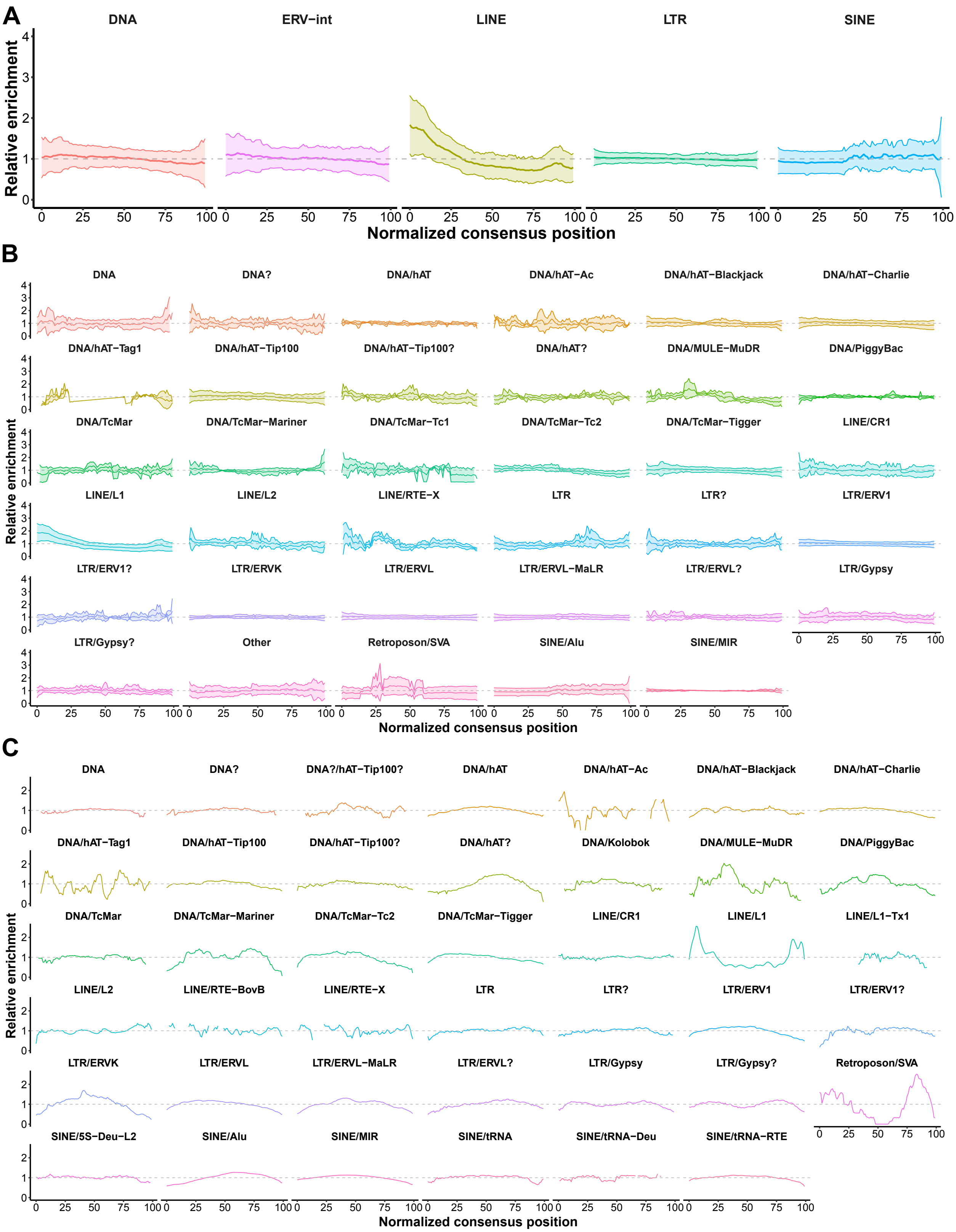
A
DNA
ERV−int
LINE
LTR
SINE
Relative enrichment
Normalized consensus position
B
DNA
DNA?
DNA/hAT
DNA/hAT−Ac
DNA/hAT−Blackjack
DNA/hAT−Charlie
DNA/hAT−Tag1
DNA/hAT−Tip100
DNA/hAT−Tip100?
DNA/hAT?
DNA/MULE−MuDR
DNA/PiggyBac
DNA/TcMar
DNA/TcMar−Mariner
DNA/TcMar−Tc1
DNA/TcMar−Tc2
DNA/TcMar−Tigger
LINE/CR1
LINE/L1
LINE/L2
LINE/RTE−X
LTR
LTR?
LTR/ERV1
LTR/ERV1?
LTR/ERVK
LTR/ERVL
LTR/ERVL−MaLR
LTR/ERVL?
LTR/Gypsy
LTR/Gypsy?
Other
Retroposon/SVA
SINE/Alu
SINE/MIR
Relative enrichment
Normalized consensus position
C
DNA
DNA?
DNA?/hAT−Tip100?
DNA/hAT
DNA/hAT−Ac
DNA/hAT−Blackjack
DNA/hAT−Charlie
DNA/hAT−Tag1
DNA/hAT−Tip100
DNA/hAT−Tip100?
DNA/hAT?
DNA/Kolobok
DNA/MULE−MuDR
DNA/PiggyBac
DNA/TcMar
DNA/TcMar−Mariner
DNA/TcMar−Tc2
DNA/TcMar−Tigger
LINE/CR1
LINE/L1
LINE/L1−Tx1
LINE/L2
LINE/RTE−BovB
LINE/RTE−X
LTR
LTR?
LTR/ERV1
LTR/ERV1?
LTR/ERVK
LTR/ERVL
LTR/ERVL−MaLR
LTR/ERVL?
LTR/Gypsy
LTR/Gypsy?
Retroposon/SVA
SINE/5S−Deu−L2
SINE/Alu
SINE/MIR
SINE/tRNA
SINE/tRNA−Deu
SINE/tRNA−RTE
Relative enrichment
Normalized consensus position

**Extended Data Fig. 4**: LINE 5' end is enriched for cCRE annotation. A) Relative enrichment of consensus sequence coverage for cCRE overlapping (foreground) vs. non-cCRE overlapping (background) elements per TE subfamily grouped by TE class. Background elements were randomly chosen while controlling for length. Mean relative enrichment across TE subfamilies +/- one standard deviation is shown. B) Same as A except TE subfamilies are grouped by TE family. C) Relative enrichment of consensus sequence coverage for region of cCRE overlap vs. entire foreground elements per TE subfamily grouped by TE family.

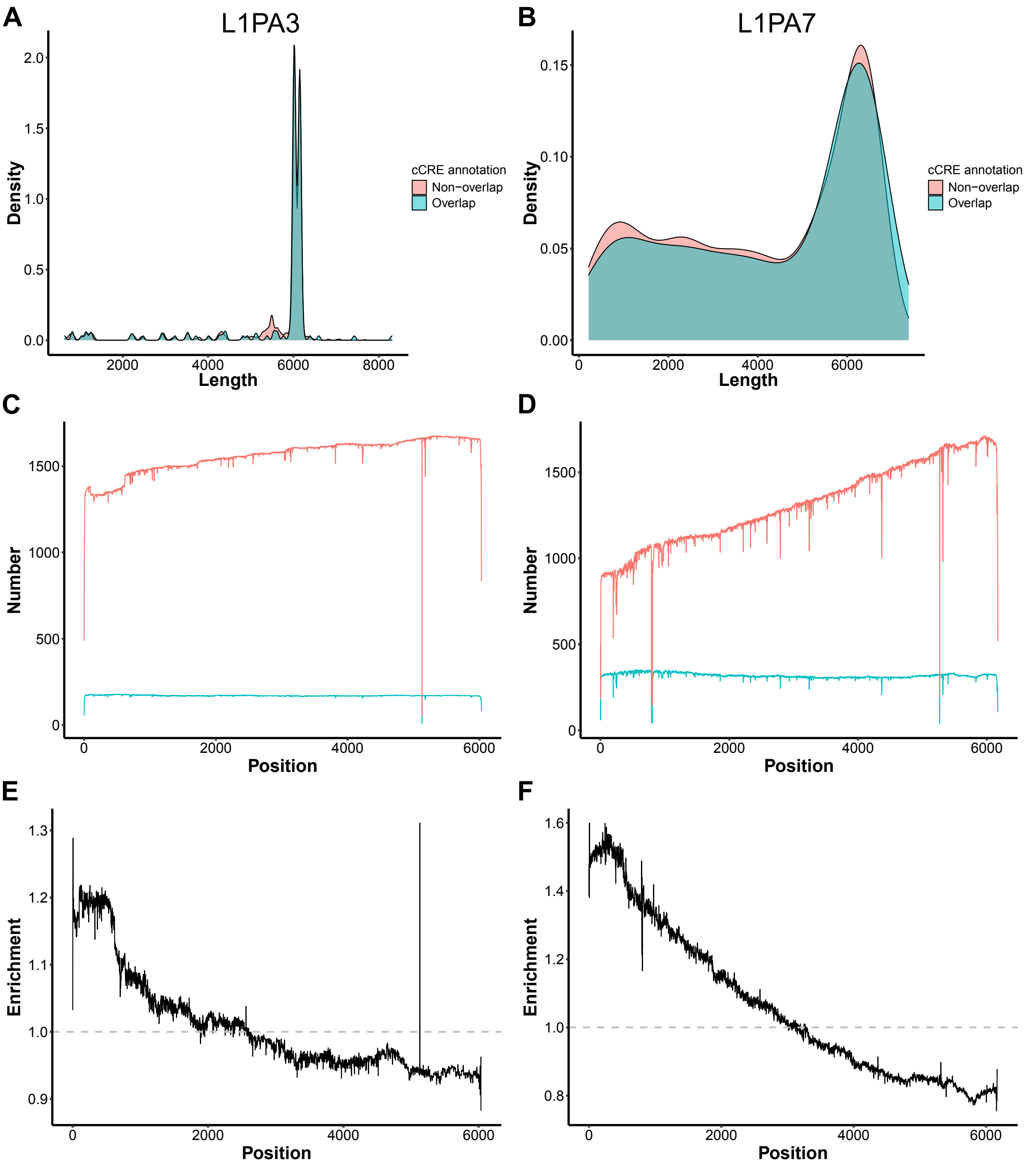


**Extended Data Fig. 5**: Examples of LINE1 subfamilies displaying 5' bias in cCREs. A) and B) L1PA3 and L1PA7 length distribution in cCRE overlapping (foreground) and non-cCRE overlapping (background) elements. C) and D) Number of elements that align across the length of the subfamily consensus sequence in foreground and background in L1PA3 and L1PA7. E) and F) Enrichment of TE subfamily consensus coverage in foreground over background in L1PA3 and L1PA7.

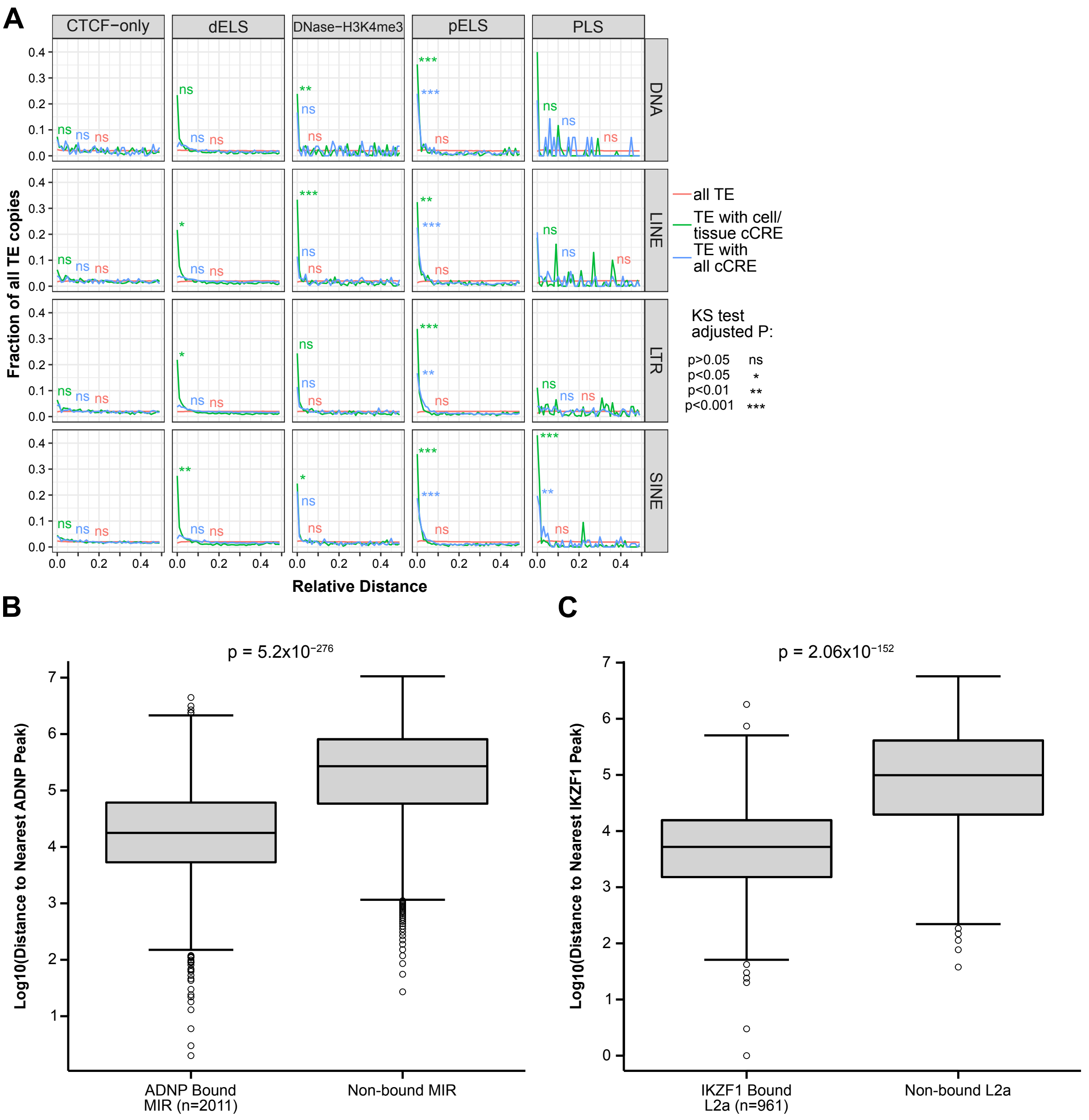


**Extended Data Fig. 6**: Distance of TEs contributing to cCREs or TFBS to non-TE sites compared to non-cCRE or non-bound TEs. A) Relative distance of all TEs to mouse cell agnostic cCREs (red), cCRE associated TEs to cell agnostic cCREs (blue), and cCRE associated TEs to same cell/tissue type cCREs (green). B) Distribution of distances from ADNP bound and non-bound MIR to their nearest non-TE ADNP binding site. C) Distribution of distances from IKZF1 bound and non-bound L2a to their nearest non-TE IKZF1 binding site. Two-sided Wilcoxon rank sum test p-values are shown.

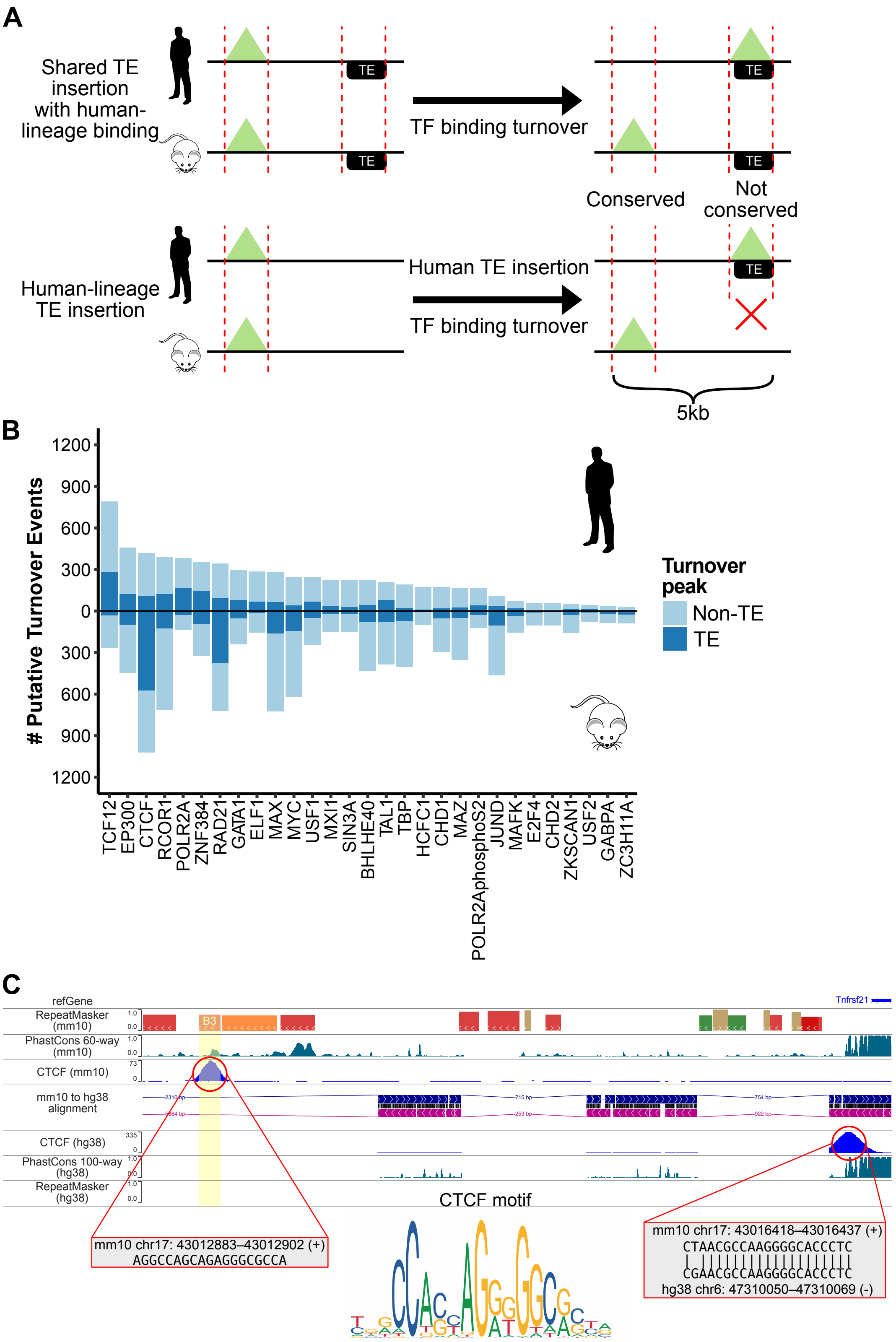
A
Shared TE insertion with human-lineage binding
TE
TF binding turnover
Conserved
Not conserved
Human-lineage TE insertion
Human TE insertion
TF binding turnover
5kb
B
### Putative Turnover Events
1200
900
600
300
0
300
600
900
1200
Turnover peak
Non-TE
TE
TCF12
EP300
CTCF
RCOR1
POLR2A
ZNF384
RAD21
GATA1
ELF1
MAX
MYC
USF1
MXI1
SIN3A
BHLHE40
TAL1
TBP
HCFC1
CHD1
MAZ
POLR2AphosphoS2
JUND
MAFK
E2F4
CHD2
ZKSCAN1
USF2
GABPA
ZC3H11A
C
refGene
RepeatMasker (mm10)
PhastCons 60-way (mm10)
CTCF (mm10)
mm10 to hg38 alignment
CTCF (hg38)
PhastCons 100-way (hg38)
RepeatMasker (hg38)
Tnfrsf21
B3
CTCF motif
mm10 chr17: 43012883–43012902 (+)
AGGCCAGCAGAGGGCGCCA
mm10 chr17: 43016418–43016437 (+)
CTAACGCCAAGGGGCACCCTC
CGAACGCCAAGGGGCACCCTC
hg38 chr6: 47310050–47310069 (-)

**Extended Data Fig. 7**: TF turnover facilitated by TEs. A) Schematic for defining putative TE-derived TF turnover events using human as an example. Two scenarios are depicted for when the TE is shared between human and mouse and when the TE is lineage specific. Only TF binding sites within 5kb of each other across syntenic regions are considered. The likely ancestral TF binding site is inferred based on conservation using phastCons score. B) Number of putative TFBS turnover events per TF in human and mouse. Each bar is split by TE or non-TE sequence origin of the new TFBS peak. C) Browser shot of CTCF binding site turnover in mouse facilitated by rodent lineage insertion of B3. Underlying CTCF motif sequence alignment in human and mouse are shown (if available).

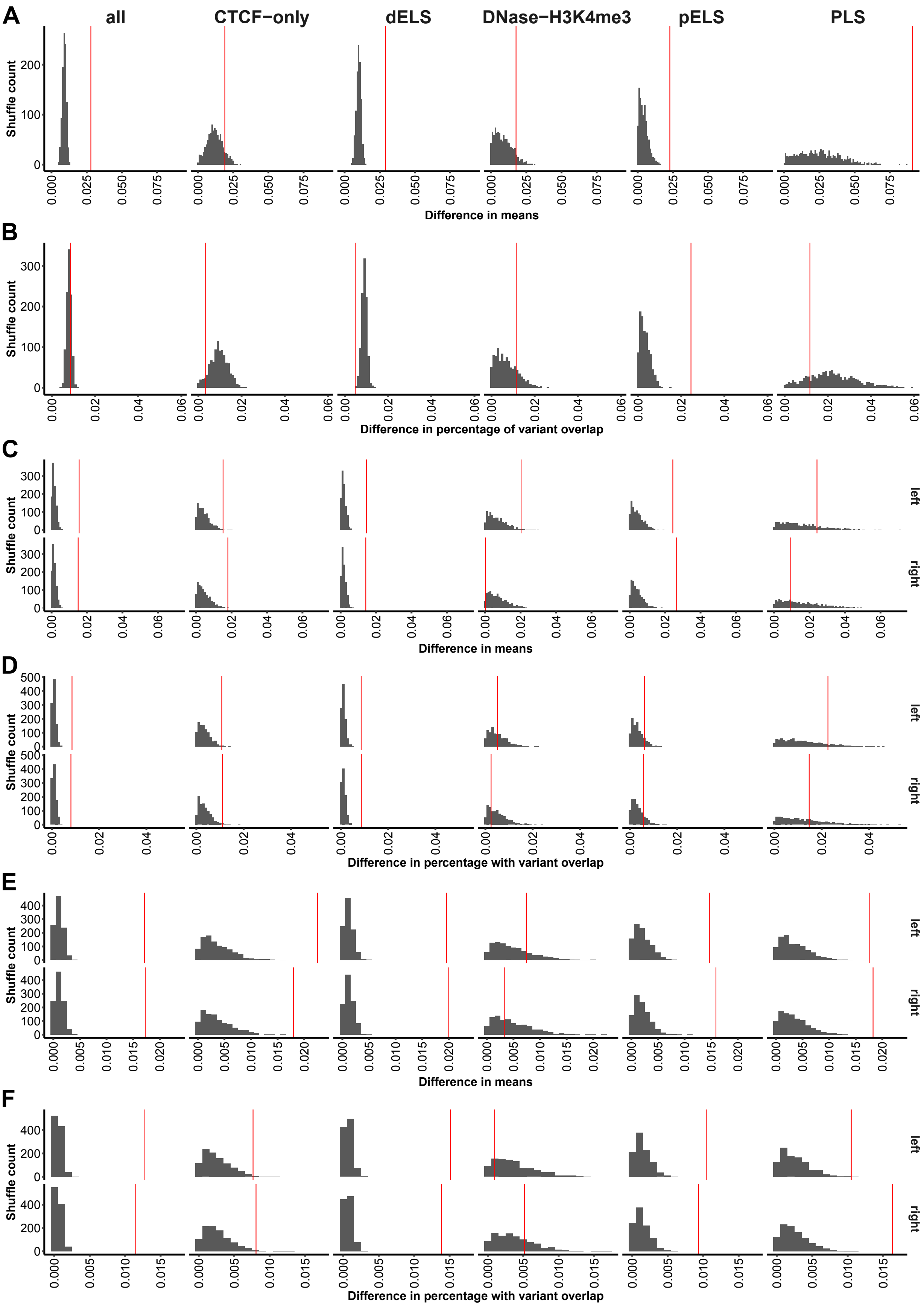

A
B
C
D
E
F
all
CTCF−only
dELS
DNase−H3K4me3
pELS
PLS
Shuffle count
Difference in means
Difference in percentage of variant overlap
Difference in means
Difference in percentage with variant overlap
Difference in means
Difference in percentage with variant overlap
left
right

**Extended Data Fig. 8**: Permutation tests comparing cCRE observations to random genomic background. Distributions of 1000 permutations are shown for A) difference in mean variants per 100bp between annotated TE-derived and non-TE cCREs, B) difference in percentage of cCREs that overlap variants between annotated TE-derived and non-TE cCREs, C) difference in mean variants per 100bp between annotated TE-derived cCREs and their flanking regions that overlap at least one variant, D) difference in percentage of cCREs that overlap variants between annotated TE-derived cCREs and their flanking regions, E) difference in mean variants per 100bp between annotated non-TE cCREs and their flanking regions that overlap at least one variant, and F) difference in percentage of cCREs that overlap variants between non-TE cCREs and their flanking regions. The observed values for cCREs is indicated by the red vertical line.

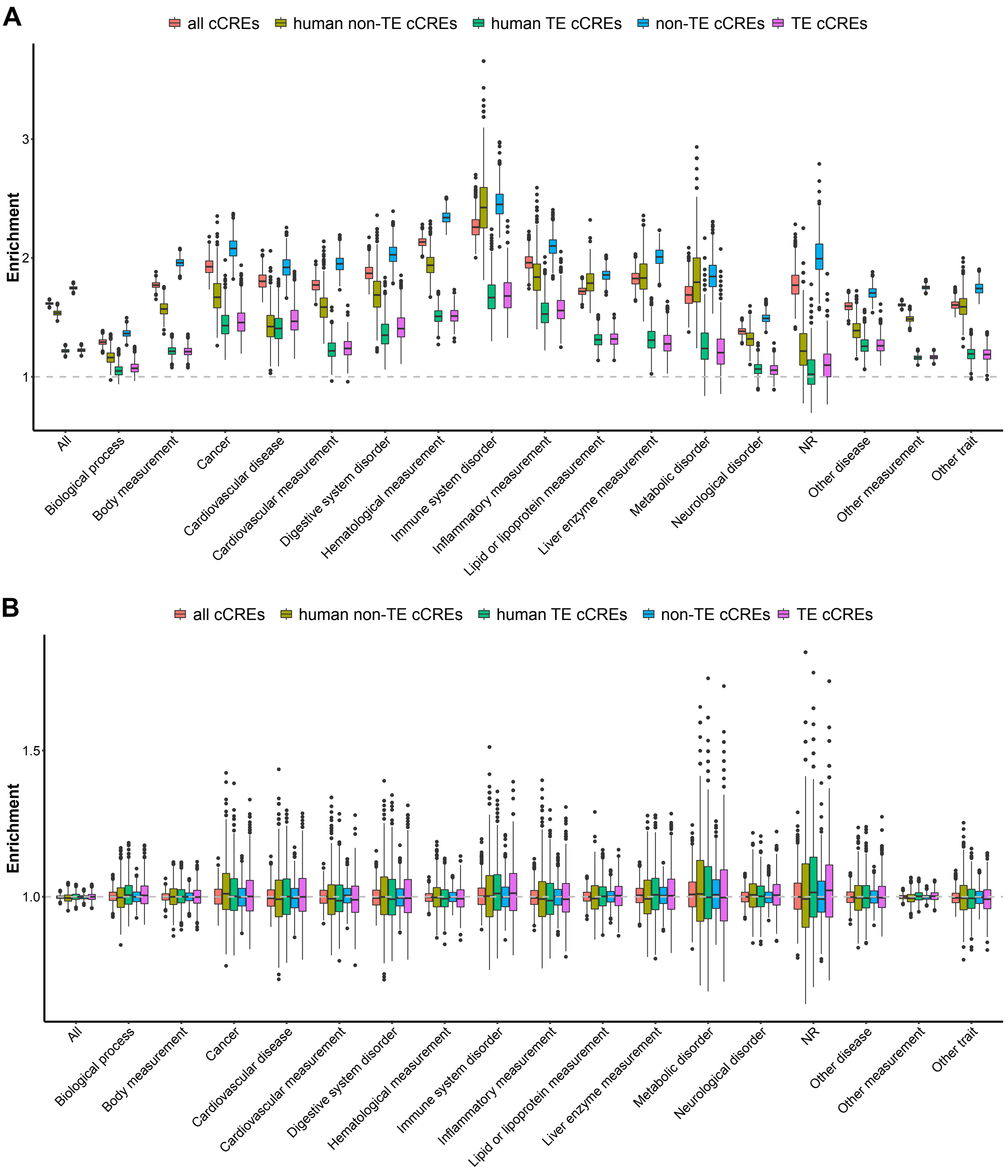


**Extended Data Fig. 9**: Enrichment of cCRE overlap with GWAS variants. A) Enrichment distribution for observed cCRE coordinates and TE annotations. B) Enrichment distribution for shuffled cCRE coordinates.
