## Supplementary Methods for "Regulatory Transposable Elements in the Encyclopedia of DNA Elements"

**TE annotations from RepeatMasker**

Repeat annotations in hg38 (Dec 2013) using RepeatMasker open-4.0.5 and Repeat Library 20140131 were downloaded from <https://repeatmasker.org/>^1^. Simple repeats, satellites, low-complexity, and unknown class repeats were removed to leave only TEs. Split TE annotations were merged using a custom python script that merges nearby TE annotations of the same family if they meet all of the following criteria: 1) they are on the same strand, 2) their positions in the corresponding TE family consensus sequence do not overlap and, when combined, match the expectation of a full-length TE annotation, and 3) the distance between them is smaller than the corresponding gap in consensus sequence.

**Human-mouse orthologous TE and syntenic cCRE classification**

We classified each cCRE starting from human or mouse based on synteny and cCRE and TE overlap (Extended Data Fig. 2A). For cCREs without synteny, they were classified as species-specific or species-TE, depending on TE overlap. For cCREs with synteny, those that overlap the same TE, based on TE family, in both human and mouse were classified as orthologous TEs. Otherwise, the cCREs were classified as syntenic.

For cCRE comparison, cCREs only found in one species, regardless of synteny annotation, were classified as species-specific cCREs. For cCREs with syntenic cCREs, they were classified as shared or different based on whether the cCRE types are the same or different between human and mouse species.

**TE subfamily consensus sequences**

We attempted to obtain consensus sequences for each TE subfamily listed in hg38 RepeatMasker annotations to investigate whether the ancestral TE sequences have TF binding motifs. For LINEs, we constructed full length consensus sequences based on the most common LINE fragments (e.g. “L1HS_5end”, “L1P1_orf2”, “L1HS_3end”) that align to each annotated LINE TE. For other TEs, we chose the most frequently aligned consensus sequence from RepeatMasker alignments in the TE subfamily if the frequency was greater than 50% and took the consensus sequence from RepBase Repeat Library RepeatMasker Edition 20170127^2^. We were unable to obtain consensus sequences for four TE subfamilies: Alu, L2d, L2d2, and MLT1B-int.

**Percent ancestral origin relationship with Kimura divergence**

Linear relationship between percent ancestral origin and Kimura percent divergence was plotted using stat_smooth with “lm” method, and r-squared was calculated using stat_cor in R.

**LINE1 5’ enrichment in cCREs**

To calculate the relative enrichment of consensus sequence coverage for cCRE associated TEs (foreground) over non-cCRE TEs (background), TE copies were first aligned to their subfamily consensus sequence. Then, the number of TE copies that cover each base of the consensus sequence was obtained for both cCRE and non-cCRE TEs. Enrichment was calculated by dividing the percentage of cCRE TE consensus coverage by the percentage of non-cCRE TE consensus coverage. In order to combine consensus enrichment across TE subfamilies, consensus sequence length was normalized to a total normalized length of 100. Enrichments were combined for each normalized length “bin” by taking the mean enrichment.

For relative enrichment of cCRE overlapping regions over entire foreground TEs, the cCRE overlapping regions of TE copies were obtained based on alignment of the whole TE copy to their subfamily consensus sequence. For each TE subfamily, the relative coverage at each consensus sequence base, defined as the cCRE overlap consensus coverage relative to foreground element consensus coverage, was calculated as the following:

$$\frac{(cCRE overlap coverage at base i)/(total number of overlapping cCREs)}{(foreground coverage at base i)/(total number of foreground)}$$

Relative enrichment was then calculated as the relative coverage divided by the mean relative coverage followed by the same binning and length normalization described above for combining TE subfamilies into TE families.

**Processing K562 lentiMPRA data**

We downloaded mean log2FC values for lentiMPRA in K562 from the ENCODE portal^3^ (<https://www.encodeproject.org/>). All log2FC values were normalized to the median of shuffled sequences, which we defined as negative controls. Tested elements in lentiMPRA were connected to their provided genomic coordinates.

**Human population variant frequency permutation tests**

Genomic regions directly flanking cCREs and random genomic regions with their flanking regions were needed to calculate the likelihood of getting percentages of variant overlap and variant frequency rates equal to or more extreme than observed cCREs. Flanking regions of the same length as the cCRE were obtained by BEDTools flank^4^. Random genomic regions and their flanks were obtained by shuffling TE-derived and non-TE cCRE coordinates within non-gapped, non-coding genome regions (as defined by GENCODEv41^5^) 1000 times using BEDTools shuffle followed by BEDTools flank^4^. We excluded any region that overlaps coding sequence from further analysis due to likely different selective pressure within coding regions. Next, we overlapped observed cCRE, shuffled cCRE, and all flanking region coordinates with common variants (allele frequency > 1%) from the 1000 genomes project after liftOver from hg19 to hg38^6,7^.

To compare between TE-derived and non-TE cCREs as well as between observed cCREs and genomic background, we used two metrics: percentage with variant overlap and variant frequency per 100bp when overlapping at least one variant. We obtained the null distribution for the difference between TE-derived cCREs, non-TE cCREs, and their respective flanks with further separation by cCRE type based on shuffled cCRE coordinates and pre-shuffling annotations. This allowed us to account for differences in cCRE length and number between groups. Null distributions were acquired for six comparisons: 1) difference in mean variants per 100bp between annotated TE-derived and non-TE cCREs, 2) difference in percentage of cCREs that overlap variants between annotated TE-derived and non-TE cCREs, 3) difference in mean variants per 100bp between annotated TE-derived cCREs and their flanking regions that overlap at least one variant, 4) difference in percentage of cCREs that overlap variants between annotated TE-derived cCREs and their flanking regions, 5) difference in mean variants per 100bp between annotated non-TE cCREs and their flanking regions that overlap at least one variant, and 6) difference in percentage of cCREs that overlap variants between non-TE cCREs and their flanking regions. Empirical p-values were calculated for differences in mean variants per 100bp and in percentage of cCREs that overlap variants compared to flanking regions (TE-derived cCRE vs. non-TE cCRE, TE-derived cCRE vs. random, and non-TE cCRE vs. random) as the probability of getting an absolute difference as large or larger than the observed difference (listed in Supplementary Table 3). For TE-derived and non-TE cCRE comparison with random genomic background, the mean empirical p-value of cCRE flanking regions (left and right) was used.

**TE overlap in genotyping arrays**

Hg19 coordinates corresponding to probed variants were downloaded for nine genotyping arrays from a previous genotyping array comparison study^8^. RepeatMasker annotations in hg19 were downloaded (RepeatMasker open-4.0.5 and Repeat Library 20140131) and intersected with genotyping array SNP coordinates using BedTools^1,4^. The number and percentage of probed variants overlapping repetitive elements was calculated based on the intersection.

**References**

1.	Smit, A., Hubley, R. & Green, P. RepeatMasker Open-4.0. *2013-2015 <http://www.repeatmasker.org>*.

2.	Bao, W., Kojima, K. K. & Kohany, O. Repbase Update, a database of repetitive elements in eukaryotic genomes. *Mob. DNA* **6**, 1–6 (2015).

3.	Agarwal, V. *et al.* Massively parallel characterization of transcriptional regulatory elements in three diverse human cell types. *bioRxiv* 2023.03.05.531189 (2023) doi:10.1101/2023.03.05.531189.

4.	Quinlan, A. R. & Hall, I. M. BEDTools: a flexible suite of utilities for comparing genomic features. *Bioinformatics* **26**, 841–842 (2010).

5.	Frankish, A. *et al.* GENCODE 2021. *Nucleic Acids Res.* **49**, D916–D923 (2021).

6.	The 1000 Genomes Project Consortium. A global reference for human genetic variation. *Nature* **526**, 68–74 (2015).

7.	Kuhn, R. M., Haussler, D. & James Kent, W. The UCSC genome browser and associated tools. *Brief. Bioinform.* **14**, 144–161 (2013).

8.	Verlouw, J. A. M. *et al.* A comparison of genotyping arrays. *Eur. J. Hum. Genet. 2021 2911* **29**, 1611–1624 (2021).
